# Leveraging single cell multiomic analyses to identify factors that drive human chondrocyte cell fate

**DOI:** 10.1101/2024.06.12.598666

**Authors:** Divya Venkatasubramanian, Gayani Senevirathne, Terence D. Capellini, April M. Craft

**Affiliations:** Department of Molecular and Cellular Biology, Harvard University, Cambridge, United States; Department of Orthopedic Research, Boston Children’s Hospital, Boston, United States; Department of Orthopedic Surgery, Harvard Medical School, Boston, United States; Human Evolutionary Biology, Harvard University, Cambridge, United States; Broad Institute of MIT and Harvard, Cambridge, United States; Harvard Stem Cell Institute, Cambridge, United States

## Abstract

Cartilage plays a crucial role in skeletal development and function, and abnormal development contributes to genetic and age-related skeletal disease. To better understand how human cartilage develops *in vivo*, we jointly profiled the transcriptome and open chromatin regions in individual nuclei recovered from distal femurs at 2 fetal timepoints. We used these multiomic data to identify transcription factors expressed in distinct chondrocyte subtypes, link accessible regulatory elements with gene expression, and predict transcription factor-based regulatory networks that are important for growth plate or epiphyseal chondrocyte differentiation. We developed a human pluripotent stem cell platform for interrogating the function of predicted transcription factors during chondrocyte differentiation and used it to test *NFATC2*. We expect new regulatory networks we uncovered using multiomic data to be important for promoting cartilage health and treating disease, and our platform to be a useful tool for studying cartilage development *in vitro*.

**Statement of Significance:** The identity and integrity of the articular cartilage lining our joints are crucial to pain-free activities of daily living. Here we identified a gene regulatory landscape of human chondrogenesis at single cell resolution, which is expected to open new avenues of research aimed at mitigating cartilage diseases that affect hundreds of millions of individuals world-wide.

## Introduction

During human skeletal development, chondrocytes participate in the patterning, growth, and function of skeletal elements^1^. Growth plate chondrocytes promote longitudinal growth, while articular cartilage chondrocytes protect joint surfaces^2,3^. Specification of chondrocytes in these morphologically and functionally distinct cartilage tissues occurs *in utero*^4–6^. Human diseases and *in vivo* animal models helped identify many genes that participate in chondrogenesis, including those encoding transcription factors^7–11^, signaling pathway components^12,13^, and extracellular matrix proteins^14^. However, many genes involved in chondrogenesis remain to be discovered.

Transcriptomic methods using *in vivo* tissues and *in vitro* models^15–19^ identified additional genes that are important during chondrogenesis. However, transcriptomes cannot fully describe how chondrocyte differentiation is regulated^18^. Single cell or single nucleus methods which can simultaneously generate single nucleus RNA transcriptomes (i.e., snRNA-seq) and single nucleus chromatin accessibility (i.e. snATAC-seq) data, referred to in this paper as multiomics, offer a way to identify important genes and the regulatory networks in which they participate. Access to gestational donor tissue and challenges in extracting single cells or nuclei from tissues with dense extracellular matrix, such as cartilage, has limited the application of multiomics to chondrogenesis.

Here we report multiomic data obtained from single nuclei recovered from developing distal femurs of a 59-day-old fetus and a 72-day-old fetus. We computationally interrogated these multiomic data individually to define chondrocyte subtypes present at these timepoints. We then integrated the snRNA-seq and snATAC-seq data at each timepoint to predict transcription factors and regulatory networks that distinguish articular and growth plate chondrocyte lineages and the different cell types within each lineage. We also developed a human pluripotent stem cell platform by which gene function could be tested during human chondrogenesis *in vitro*.

## Results

### Combining single nucleus transcriptome and open chromatin data predicts different cell types in developing fetal distal femur

The cartilage long bone anlage during fetal development contains chondrogenic cells in different states of differentiation and maturation^2,3^. The anlage includes cells that will become part of the permanent articular cartilage and cells that will function in growth plates. We obtained distal femurs from two fetal donors at embryonic day 59 (E59) and 72 (E72). Chondrocyte precursors to articular chondrocytes and growth plate chondrocytes were present in the distal femur at both timepoints based on histologic assessment of stage matched tissues (**Figure 1A**). The E72 specimen, being later in development and larger in size compared to the E59 specimen, was expected to have more cells and altered proportions of differentiated cells (e.g., E72 had a more apparent hypertrophic chondrocyte zone, **Figure 1A**). In a recent effort to define the Carnegie stages of human embryos, it was found that E59 corresponds to Carnegie stage 23^20^. By the late E60 and E70 timepoints, it was proposed that soft tissues, such as the anterior/posterior cruciate ligaments and synovium, emerge to tether the distal femur to the proximal tibia, suggesting that additional connective tissue-like cells could be captured in the E72 specimen. Nuclei were recovered from each specimen and subjected to snRNA-seq and snATAC-seq. Shared barcodes from this joint multiomic profiling allowed us to assign unique RNA-seq and ATAC-seq profiles to each nucleus. We obtained profiles for 5,056 nuclei from the E59 specimen and 13,865 nuclei from the E72 specimen.

**Figure 1.**
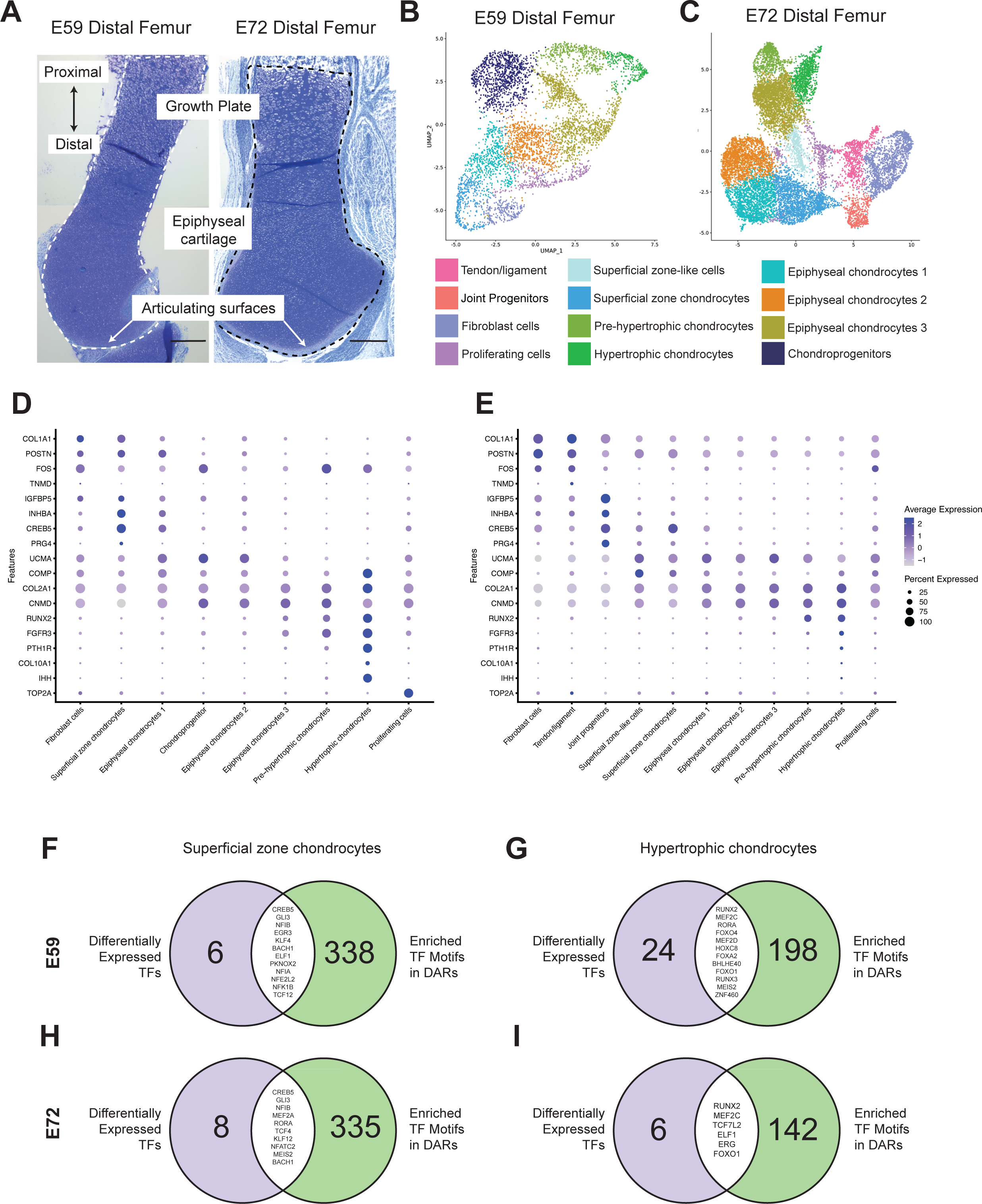
Transcriptomic and epigenetic landscape of chondrocyte subtypes within the developing human distal femur. A. Histological sections of distal femur cartilage stained with toluidine blue on embryonic day (E)59 and E72. Proximal/distal orientation, and general location of phenotypically distinct chondrocyte subtypes (i.e., those at the articulating surface, within the epiphysis and comprising the growth plate) are noted. Cells were isolated from the area indicated in dotted lines. Scale bar, 500 µm. (B-C) UMAP projection and wKNN clustering of cells isolated from the distal femur at E59 (B) and E72 (C-(D-E) Dot-plots represent normalized Log(2)FC expression of marker genes and the percentage of cells expressing each marker at E59 (D) and E72 (E). (F-I) Venn diagrams showing intersection of differentially expressed TFs (purple, left) and differential TF motif accessibility (green, right) in each representative cluster at E59 (F,G) and E72 (H,I). The overlap represents TFs that are both differentially expressed and whose motifs are differentially accessible in the same population.

To infer the cell types from which these nuclei came, we performed unsupervised clustering using a weighted nearest neighbor (wKNN) analysis^21^, which integrates transcriptomic and open chromatin data by weighting their relative utility in each cell. This led us to identify 10 cell type clusters at E59 and 16 cell type clusters at E72. We computationally removed hematopoietic (*CD34*^high^)^22^, skeletal muscle (*MYF5*^high^)^23^, and neuronal (*GFAP*^high^)^24^ cells (**Supplemental Table S1**), re-clustered the remaining cells, and named the clusters based on their patterns of differentially expressed genes (DEGs) (**Figure 1B-C, Figure S1, Supplemental Tables 2 and 3**). Superficial zone-like chondrocytes (SZC) were enriched for *CREB5*, *FAP*, and *INHBA* expression, with a subset also expressing *PRG4*. Hypertrophic chondrocytes (HC) and pre-hypertrophic chondrocytes (PHC) were enriched for *RUNX2*, *FGFR3*, and *PTH1R* expression with the HC cluster also expressing *COL10A1*, *IHH,* and *PANX3* (**Figures 1D-E, S1**).

Chondrocytes with patterns of DEGs that differed from those associated with articular chondrocytes (SZC) or growth plate chondrocytes (HC) comprised ∼ 50% of the cells in E59 and E72 femurs. We named these “Epiphyseal” chondrocytes because all expressed cartilage associated proteins (e.g., *COL2A1*, *COL9A1*, *ACAN, COMP*) and transcription factors (e.g., *SOX9*, *SOX5*, and *SOX6*)^25^, and further subdivided them into 3 clusters (EC1, EC2, EC3). In UMAP space EC1 was closer to the SZC cluster and expressed *UCMA* and *COMP* (**Figure 1B-E**). EC3 was closer to the PHC and HC clusters and had higher expression of *FGFR3* and *BMPR1B* than other epiphyseal clusters. Other clusters in our datasets likely represented non-chondrogenic connective tissue and progenitor cells based on their unique DEGs, and included fibroblast-like cells (*COL1A1*)^26^, found at E59 and E72, and joint progenitor cells (*COL1A1*, *IGFBP5*) and tendon/ligament cells (*TNMD)*^27^ retrieved at E72 (**Figure 1D-E, S1C-D**).

### Single nucleus chromatin accessibility uncovered cluster-specific enrichment of transcription factor binding motifs

Among all recovered nuclei, snATAC-seq found 152,574 open chromatin regions at E59 and 178,116 open chromatin regions at E72. Overlapping regions in the 2 samples numbered 124,072. Using the transcriptomic-based naming strategy of cell type clusters described above, we compared open chromatin regions between cell types, searching for differences. At E59 there were 42,693 differentially accessible regions (DARs) between cell types, and there were 88,056 DARs at E72 (**Supplemental Tables 4 and 5)**. Consistent with chromatin accessibility being associated with gene expression, SZCs were enriched for open chromatin near *PRG4* (e.g., chr1:186,292,811-186,293,287), and these peaks were also detected in EC1, even though *PRG4* is not expressed in this cell type (**Figure S2A,C)**. Similarly, HCs were enriched for open chromatin near *COL10A1* (e.g., chr6:116,144,312-116,144,927; **Figure S2E,G**), as were PHCs and EC3 even though these cells were not expressing *COL10A1*.

Because transcription factor (TF) binding to open chromatin can control gene expression, we used a TF consensus binding site search algorithm to predict TFs that may be active in different cell clusters^28^. We found the algorithm detected signatures for TFs with known cell type-specific functions in cartilage, such as *RUNX2* in HCs and *CREB5* in SZCs (**Supplemental Tables 6 and 7**)^29,30^. The HC cluster contained cells for which snRNA-seq indicated *RUNX2* expression, snATAC-seq detected enriched accessibility of the RUNX2 consensus binding sequence (motif), and snRNAseq detected expression of a known RUNX2 target gene *COL10A1* (**Figure S2D, H**). Similarly, the SZC cluster contained cells for which snRNA-seq indicated *CREB5* expression, snATAC-seq detected enriched accessibility of the CREB5 consensus motif, and snRNA-seq detected expression of a known CREB5 target gene *PRG4* (**Figure S2B,F**).

Having detected TFs with known roles in differentiated chondrocytes, we next sought to identify TFs with previously unappreciated roles in cartilage development. We did this by identifying TFs that represented DEGs between cell types using snRNA-seq and asked which TFs had motifs that were enriched in open chromatin regions identified using snATAC-seq. The number of DE TFs, the number of TF binding motifs that were differentially active, and the intersection between the two are shown in **Figure 1F-I** for representative SZC and HC clusters. TFs that fell into both categories included TFs with known roles in chondrogenesis^10,29,31^ and TFs that may have important roles in chondrogenesis (**Figure 1F-I**; **Table 1**). TFs found uniquely at either timepoint could suggest diminishing or emerging cell types at different stages of development, or differential usage of TF networks. Reassuringly, TFs that are known to be critical in specific cell types (e.g. *CREB5*, *RUNX2*) were found at both time points. We suggest that other TFs identified in this same manner are likely to have cell type-specific roles and thus warrant further investigation.

**Table 1.** TFs that are both differentially expressed and whose motif is significantly enriched in the same cluster.

| Cluster | Transcription Factor(s) |
| --- | --- |
| **E59** |  |
| Fibroblast cells | NFIB, KLF4, NFIA, RFX7, SOX4 |
| Proliferating cells | KLF3, MAFG, MAZ, PBX3, PKNOX1, SOX8 |
| Superficial zone chondrocytes | BACH1, CREB5, EGR3, ELF1, GLI3, KLF4, NFE2L2, NFIA |
| Epiphyseal chondrocytes 1 | BACH1, BACH2, CREB5, NFATC1, NFIB, NFKB1 |
| Chondroprogenitors | STAT3, FOXC1, NFIA, NFIB, RFX3 |
| Epiphyseal chondrocytes 2 | BACH2, SNAI2 |
| Epiphyseal chondrocytes 3 | ETS1, EBF1, RUNX2, RUNX3 |
| Pre-hypertrophic chondrocytes | FOXA3, FOXO1, LEF1, RUNX2, RUNX3, SOX6, TEAD4 |
| Hypertrophic chondrocytes | BHLHE40, FOXA2, FOXO1, FOXO4, HOXC8, MEF2C, MEF2D, MEIS2, RORA, RUNX2, RUNX3, ZNF460 |
| **E72** |  |
| Fibroblast cells | ATF3, ATF6, BACH1, BACH2, CEBPB, CEBPD, EBF2, EBF3, EGR1, EGR3, ELF2, FOS, HES1, IRF1, JDP2, JUN, JUNB, KLF10, KLF4, KLF6, MAFB, MAFF, MEOX2, MYC, NFE2L2, NFIA, NFIB, NFIL3, PBX1, REL, SOX11, SOX4, STAT3, TCF12, TWIST1, TWIST2 |
| Joint progenitors | ATF6, BACH1, CREB5, DLX3, GLI3, HMBOX1, KLF13, KLF3, KLF4, KLF6, MAF |
| Tendon/ligaments | none found |
| Proliferating cells | CREB3L2, EGR1, KLF12, KLF2, NFIB, RFX7, SOX6, TCF4 |
| Superficial zone-like cells | BACH1 |
| Superficial zone chondrocytes | BACH1, CREB5, GLI3, KLF12, MEF2A, MEIS2, NFATC2, NFIB, RORA, TCF4 |
| Epiphyseal chondrocytes 1 | FOXC1, HIF1A |
| Epiphyseal chondrocytes 2 | EBF1, RUNX1 |
| Epiphyseal chondrocytes 3 | ERG, RUNX1 |
| Pre-hypertrophic chondrocytes | ERG, ETV6, FOXK1, MECOM, RUNX1, RUNX2 |
| Hypertrophic chondrocytes | ELF1, ERG, FOXO1, MEF2C, RUNX2, TCF7L2 |

### Enhancer-based gene regulatory network analysis predicted new transcription factor networks putatively involved in cartilage development

Having defined cell types based upon their transcriptomes and open chromatin, we next sought to identify enhancer-based gene regulatory networks (eRegulons) by combining TF expression, TF motif enrichment, target gene region accessibility, and target gene expression using the SCENIC+ algorithm^32^. This predicted 37 positively correlated or ‘activating’ eRegulons at E59 and 94 at E72 (**Supplemental Tables 8 and 9**). These eRegulons predict a master TF, and accessible regions (potential enhancers) and target genes putatively controlled by these TFs. Although this algorithm also captured repressor networks (denoted with -_+ suffix in **Supplemental Tables 8 and 9**), we did not consider them further due to the reported high false positive rate in predicting repressor functions^32^. We found eRegulon-controlling TFs that have been previously implicated in cartilage development, cartilage- or bone-related diseases, or GWAS peaks associated with ‘body height’ or ‘osteoarthritis’ (**Supplemental Table 21**), emphasizing the human biological relevance of the data uncovered by these analyses. However, several of them have not yet been linked to skeletal or joint biology. These new eRegulons emerge as novel pathways involved in chondrogenesis, including some that may be differentially active across cell types within the distal femur or over developmental time.

After validating that SCENIC+ identified well characterized TFs and predicted known targets of these TFs (**Supplemental Tables 8 and 9**), we used these data to explore eRegulons that were shared across both timepoints. Cells were re-clustered using dimensionality reduction based on eRegulon activity, independently of their original transcriptomic-based UMAP and naming strategy (**Figure 2A,B**). While the organization of cells in the eRegulon UMAP was largely similar to the organization in the wKNN UMAP, some changes in relative position of cells prompted our renaming of some clusters (compare **Figures 2A,B** to **1B,C,** and **S3A,B**). The ‘Chondroprogenitor’ cluster was renamed ‘Epiphyseal chondrocytes 2b’ at E59. At E72, we merged the ‘Superficial zone-like cells’ cluster and the ‘EC1’ cluster. The EC2 and EC3 clusters at E72 also showed significant overlap. We thus considered these to be one cell population which we refer to as EC3. Finally, the majority of ‘Proliferating cells’, which exhibited *TOP2A^high^* expression, were now found interspersed within other cell clusters. The predicted activity of each eRegulon was visualized in these new UMAPs using feature plots of the level of TF expression based on scRNA-seq (**Figure 2C-H**, left columns), and their regulatory activity based on motif accessibility and expression of target genes (**Figure 2C-H**, right columns). As expected, some eRegulons were shared across E59 and E72 cells, including TFs not well established in cartilage development. Some genes, such as the known *SOX* trio of TFs and the newly uncovered *PEG3* were active in chondrocytes but not in fibroblasts (**Figure 2C,F**). *CREB5* was highly active in SZCs and EC1 clusters, along *NFE2L2*, *DLX3*, and *NFIB* (**Figure 2D,G**). We also uncovered *NR3C1*, *ZEB1*, and *TCF7L2* acting in HC, PHC, and EC3 clusters, along with known growth plate gene *MEF2C*, suggesting a role for these TFs in growth plate lineage specification (**Figure 2E,H**).

**Figure 2.**
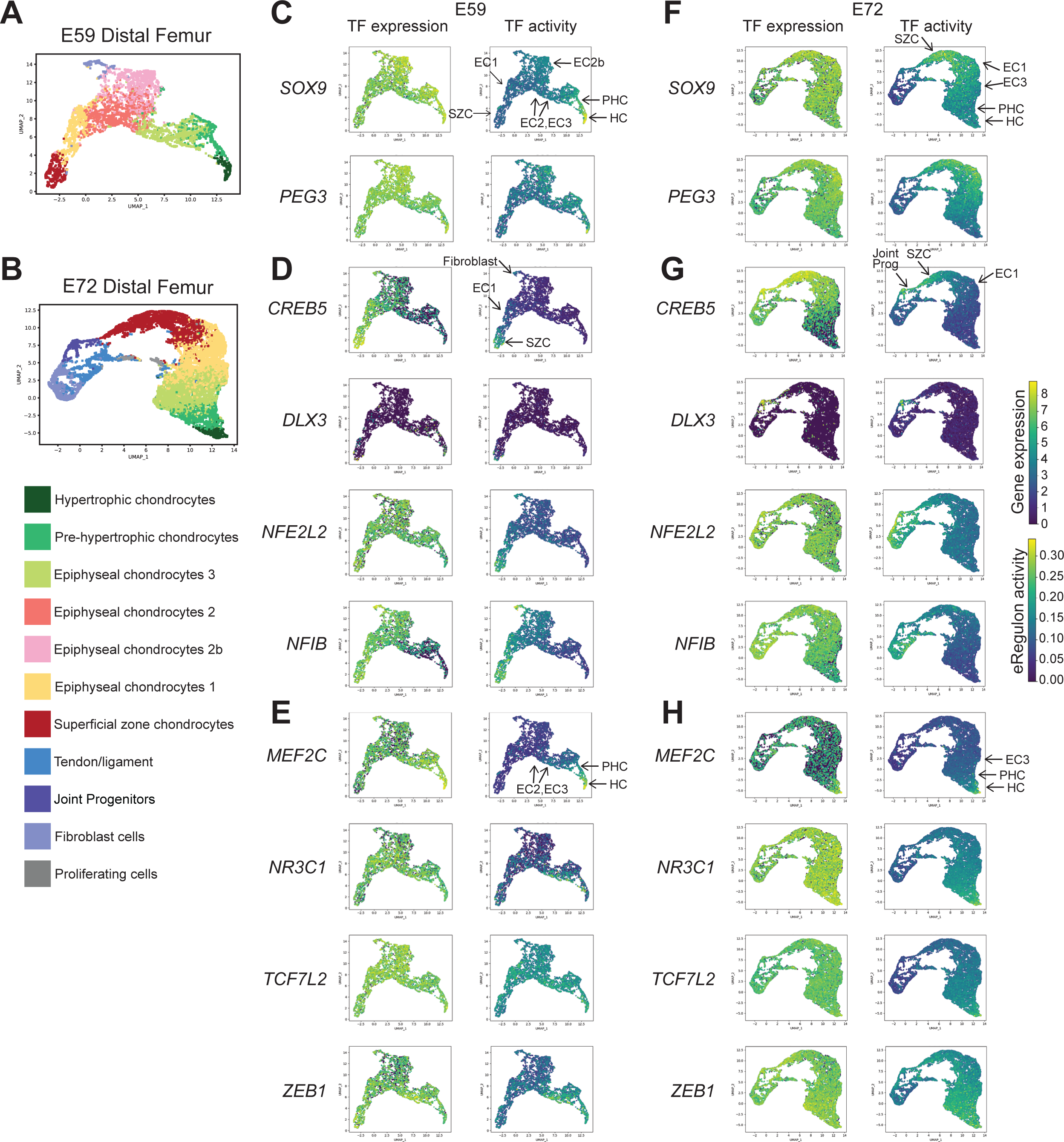
Activity of representative eRegulons (TFs) identified at both E59 and E72 using enhancer-based gene regulatory network analysis. (A-B) UMAP projection of cells and cell cluster assignments based on eRegulon activity at E59 (A) and E72 (B). Note, the wKNN cell cluster names projected onto this eRegulon UMAP are shown in Supplemental Figure 3A,B. (C-E) Feature plots depicting single cell-level expression (left) and activity (right) of eRegulons at E59. (F-H) Feature plots depicting single cell-level expression (left) and activity (right) of eRegulons at E72. eRegulons include those enriched in chondrocytes *SOX9* and *PEG3* (C,F); the articular chondrocyte lineage *CREB5*, *DLX3*, *NFE2L2* and *NFIB* (D,G); and the hypertrophic chondrocyte lineage *MEF2C*, *NR3C1*, *TCF7L2* and *ZEB1* (E,H). Arrows indicate location of cell clusters.

We next computationally calculated (ranked) the specificity of each eRegulon in different cell types using ‘regulon specificity scores’ (RSS), in which a highly ranked eRegulon in a cell cluster is predicted to be more active in that cluster compared to other eRegulons in the dataset (**Supplemental Tables 10 and 11**). The top ten eRegulons ranked in each cluster are illustrated in **Figures 3A, 4A** and **Figure S3C**. eRegulons can be active in more than one cluster, and the majority of the top ranked eRegulons were ranked highly in more than one cluster. These data, including the information gained by predicting their target genes (**Tables 2 and 3, Supplemental Tables 8 and 9**), enabled us to infer which TFs and TF families are regulating the cell fates found in the distal femur at these two timepoints.

**Figure 3.**
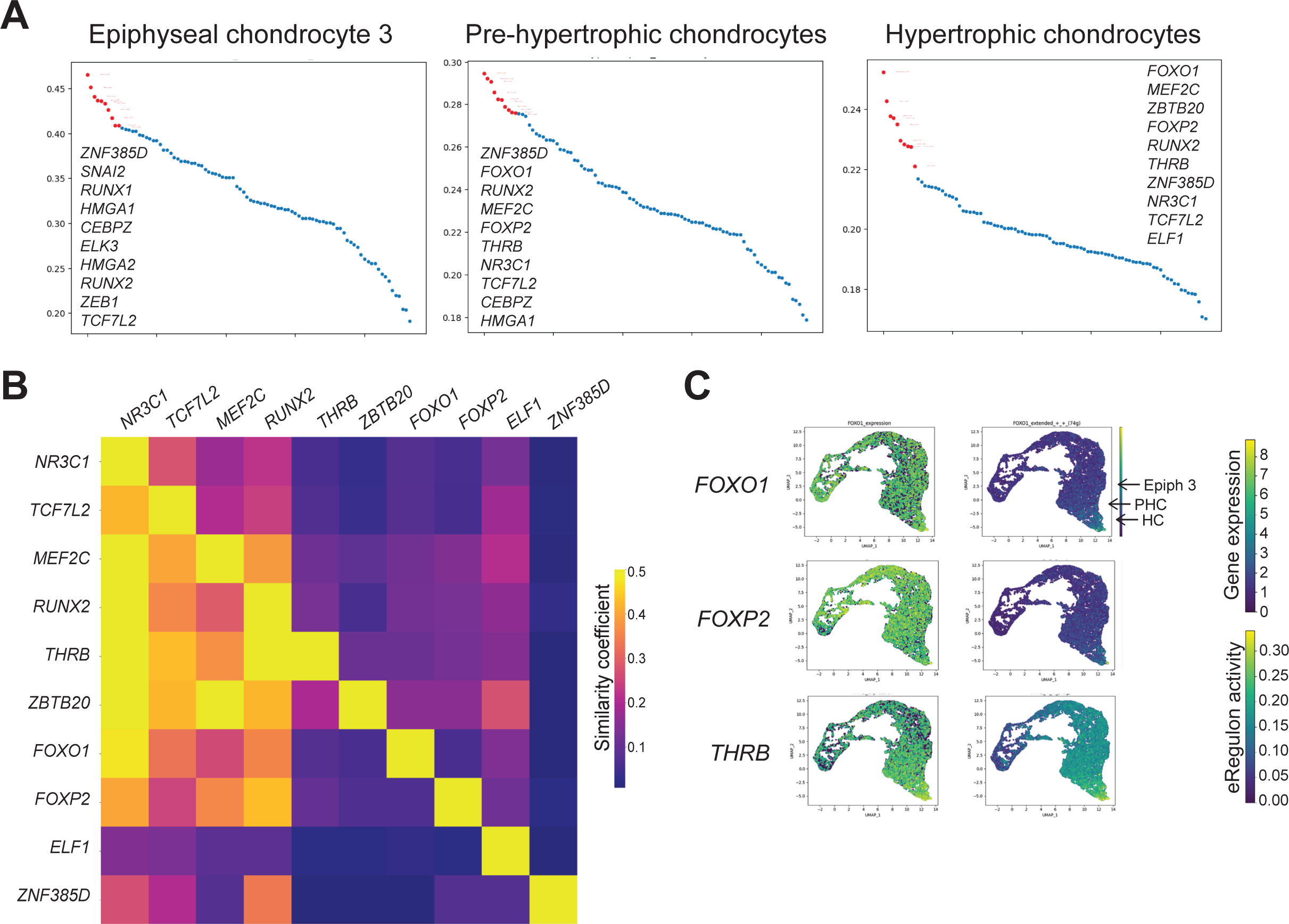
Top eRegulons in growth plate cartilage clusters at E72. (A) Scatter plot representing the top ten eRegulons, ranked by RSS, in EC3, PHC, and HC clusters. (B) Correlation heatmap representing similarity between target genes putatively controlled by the top ten eRegulons in the HC cluster. Note, differences in total target genes assigned to each TF cause heatmap to be asymmetrical. (C) Feature plots depicting single cell-level expression (left) and activity (right) of eRegulons at E72. Arrows indicate location of cell clusters.

**Table 2.** Curated list of eRegulons and targets in distal femur cartilage at E59.

| Master regulator | Tenogenic and Fibroblast genes | Articular chondrocyte genes | Epiphyseal chondrocyte genes | Growth plate chondrocyte genes |
| --- | --- | --- | --- | --- |
| CREB5 | AQP1 OSR2 COL5A3 COL5A2<br>OGN FAP SCX IGFBP5 | CREB5 SULF2 OGN HBEGF | INHBA |  |
| DLX3 |  | PRG4 |  |  |
| NFIB | COL5A2 OGN FAP COL1A2 TNC<br>IGFBP5 VCAN COL1A1 | NFIB OGN BACH1 DKK3 CREB5<br>FOXP1 KLF4 | NFIA |  |
| NFEL2L2 | FAP | TGFB3 DKK3 DCX | BARX2 BMPR1A INHBA |  |
| PLAG1 |  | FOXP1 |  |  |
| ERG |  | TGFB3 | ERG |  |
| CEBPB | EGR2 | CCN6 TIMP3 UCMA | COL2A1 SOX9 |  |
| FOXP1 |  | PTHLH LATS1 CREB5 NFKB1<br>FOXP1 | FOXC2 ERG BMPR1B SOX6<br>TGFB2 | EHD1 |
| ATF4 | COL5A1 FMOD IGFBP2 IGFBP6 | SULF2 ETV5 ELK3 RUNX1 PITX1<br>MBP PTHLH BACH1 LTBP3 NFKB1<br>KLF4 TGFB3 | FOXC2 NFATC1 NFIA CNMD<br>BARX1 | CEBPB |
| NFATC1 | IGF1R | FOXP1 NFATC1 NFIB PRG4 | ERG RFX2 NFATC1 EXT1 NFIA |  |
| SOX5 | IGF1R | IL1R1 | ERG BMPR1B SOX5 SOX6<br>BMPR1A COL11A1 TGFB2 | ENPP1 LRP6 |
| DLX5 |  | CHAD LHX9 APOD | COL2A1 | ALPL LIFR PAX3 PTH1R WNT5A<br>IHH |
| MEF2C | IGF1R | CCN6 TIMP3 CHAD | COL2A1 EXTL3 NKX3-2 | STAT1 MEF2C LIFR FGFR3<br>WNT5A SP7 EFHD1 FOXA2<br>RUNX2 PTH1R IHH PAX3<br>WNT5B RUNX3 DLX5 |
| SOX9 | UGDH | APOD CHAD FLI1 CCN6 | COL11A2 COL11A1 COL2A1<br>SOX9 COL9A1 NKX3-2 | STAT1 FGFR3 WNT5A SP7 RUNX2<br>IHH DLX5 |
| SOX6 |  |  | SOX6 |  |
| CEBPD | DCN EGR2 | CHAD DCN | COL2A1 | LIFR |
| RUNX3 | IGF1R | CHAD TIMP2 IL1R1 | COL2A1 COL9A1 NKX3-2 | ATOH8 MEF2C FGFR3 WNT5A<br>SP7 RUNX2 WNT5B RUNX3 DLX5 |
| RUNX2 | IGF1R MKX | CHAD SESN1 TIMP3 | NKX3-2 COL2A1 | MEF2C LIFR SNORC FGFR3<br>WNT5A SP7 RUNX2 PTH1R IHH<br>PAX3 RUNX3 DLX5 |

**Table 3.** Curated list of eRegulons and targets in distal femur cartilage at E72.

| Master regulator | Tenogenic and Fibroblast genes | Articular chondrocyte genes | Epiphyseal chondrocyte genes | Growth plate chondrocyte genes |
| --- | --- | --- | --- | --- |
| JUN | FBLN2 OGN FBLN1 POSTN COL5A1<br>EGR2 IGF1 VCAN OSR2 COL1A1<br>COL1A2 DCN COL3A1 TNMD<br>COL5A2 | KLF4 OGN SESN1 NFIB FGF18<br>NFATC3 DCN BGN | NFIA SULF1 | EHD1 |
| TWIST1 | COL1A1 COL5A1 OSR2 FBLN1 IGF1 |  |  |  |
| PRRX1 | COL1A1 TNC COL5A1 COL5A3 VCAN<br>POSTN | MEOX1 NFKB1 CILP WNT9A | SULF1 RFX2 |  |
| EBF1 | FBLN2 LOXL1 VIM IGFBP5 | ETV5 BMP7 FGF18 CHAD RUNX1 | NFIA | CEBPB |
| EBF2 | IGFBP6 VCAN | MBP |  |  |
| EBF3 | DCN OGN |  | NFIA |  |
| EGR1 | EGR2 FBLN1 |  | SULF1 |  |
| YY1 | TNC EGR2 | LTBP1 LTBP3 |  |  |
| MAFF | COL5A1 FBLN1 DCN IGF1 | DCN FGF18 | SULF1 |  |
| KLF4 | FBLN2 OGN FBLN1 IGFBP6 COL5A1<br>OSR2 COL1A1 COL1A2 DCN | KLF4 OGN TIMP2 HBEGF FGF18 DCN BGN | EXT1 | ALPL |
| CREB5 | IGFBP2 IGFBP5 FAP SCX | BACH1 CREB5 SULF2 CILP TGFβ3 NFATC3 GREM1 | INHBA BARX1 SOX5 |  |
| ELF2 | DCN | IL1R1 DCN |  | LIFR WNT5B |
| RUNX1 |  | IL1R1 BMP7 RUNX1 | EXT1 |  |
| BACH1 | FBLN2 TNC OGN AQP1 IGFBP2<br>LOXL1 COL5A3 UGDH FAP COL3A1<br>COL5A2 | KLF4 OGN BACH1 CREB5 LTBP3 SULF2 CILP FOXP1<br>TIMP2 NFKB1 HBEGF WNT9A TIMP3 LTBP1 DKK3 | EXT1 INHBA BMPR1A SULF1<br>SOX5 ACVR1 | EHD1 |
| CUX1 |  | SULF2 PRG4 CREB5 | SOX5 ACVR1 SOX6 ERG<br>BMPR1A |  |
| DLX3 |  | TIMP3 |  |  |
| NFE2L2 | FBLN2 TNC POSTN VIM COL5A3<br>IGFBP5 COL3A1 COL5A2 | BACH1 LTBP3 LATS1 FOXP1 TIMP2 NFKB1 HBEGF<br>WNT9A FGF18 EXT2 RUNX1 DKK3 BGN | EXT2 INHBA | DLX5 |
| NFATC2 |  | UCMA NFATC2 GREM1 | SOX6 BARX2 FOXC2 ERG<br>NFATC1 BARX1 |  |
| NFIB | TNC COL5A1 COL1A1 COL1A2 | CREB5 PITX1 ETV5 LEPR TGFβ3 FOXP1 NFIB<br>NFATC3 DKK3 | BMPR1A NFATC1 SULF1<br>NFIA RFX2 | DLX5 LIFR |
| TCF12 | COL5A1 EGR2 VIM COL1A1 COL3A1 | KLF4 BACH1 LTBP3 ETV5 TGFβ3 LATS1 TIMP2 MBP<br>NFATC3 EXT2 | NFATC1 SULF1 NFIA | WNT5B LIFR VEGFA |
| NFIA | EGR2 OSR2 TNMD | KLF4 BACH1 SESN1 FOXP1 NFIB NFKB1 FGF18<br>NFATC3 | BMPR1A NFIA |  |
| MEF2A |  | NFATC2 DCX GREM1 CREB5 | SOX5 COMP SOX9 |  |
| ERG | FMOD |  | SOX5 SOX6 FOXC2 ERG<br>BMPR1B BARX1 |  |
| HIC1 |  | MBP |  |  |
| CEBPD | EGR2 |  | SULF1 ACVR1B PROCR | LIFR CEBPB |
| FOXP2 |  |  |  | RUNX2 MEF2C |
| CEBPB | AQP1 VIM | IL1R1 MEOX1 CILP | ACVR1B PROCR | CEBPB WNT5B |
| TCF7L2 | IGF1R |  | ERG COL2A1 TGFβ2 BMPR1B<br>COL11A1 | WNT5A MATN1 ENPP1 FOXA2<br>IHH FGF3 MEF2C |
| SOX5 |  | FLI1 NFATC2 COL15A1 | ERG COL11A1 SOX5 COL9A1 | ENPP1 COL10A1 |
| SOX6 |  | FLI1 NFATC2 | COL11A1 COL9A1 |  |
| SOX9 | FMOD |  | COMP SOX9 |  |
| RUNX2 |  | IL1R1 FOXP2 | COL2A1 CNMD COL11A1<br>TGFβ2 COL11A2 BMPR1B<br>NKX3-2 COL9A1 | PTH1R FGF3 MEF2C COL10A1<br>RUNX2 ENPP1 RUNX3 MATN1 |
| THRB |  |  | BMPR1B COL11A2 COL2A1<br>COL9A1 | IHH |
| NR3C1 | MKX | IL1R1 CHAD | ERG COL2A1 TGFβ2 BMPR1B<br>COL11A2 SOX9 RFX2 | SNORC WNT5A MATN1 ENPP1<br>WNT5B RUNX3<br>IHH PTH1R MEF2C RUNX2 |
| MEF2C | MKX | CHAD<br>SESN1 | COL2A1 COMP COL11A1<br>BMPR1B NKX3-2 | SNORC PTH1R STAT1 WNT5A<br>MEF2C RUNX2 IHH MATN1 |
| ELF1 | FBLN1 IGFBP6 IGF1R VIM | SESN1 TIMP2 | BMPR1A TGFβ2 RFX2 | WNT5B LIFR COL10A1 |

Hypertrophic chondrocytes showed high eRegulon activity of *RUNX2* and *MEF2C* at both E59 and E72. The data generated by SCENIC+ also predicted that they transcriptionally activate similar target genes (**Figures 3B, S3D, S4)**, including known growth plate regulators such as *PANX3* and *IHH* at E59 and *PTH1R* and *MATN1* at E72 (**Tables 2 and 3, Supplemental Tables 8 and 9**). A subset of HC-enriched eRegulons were uncovered exclusively in one timepoint (**Figure 3B,C, Figure S3D,E**). Despite these differences, many of the top ten most specific eRegulons in the growth plate lineage at each time point show high similarity in the genes they putatively control (**Figure 3B, S3D**), suggesting cooperativity between TFs in driving growth plate gene programs. Interestingly, these analyses predicted that some TFs regulate the expression of other master TFs, pointing to a potential hierarchy of regulation in driving growth plate specification (**Supplemental Tables 8 and 9**).

As less is understood about the specification of epiphyseal chondrocytes into articular chondrocytes, we next focused on these lineages. We had already found that SZCs at both timepoints showed high activity of the *CREB5* eRegulon, which putatively controls articular cartilage genes such as *SULF2* and *FGF18 (***Tables 2 and 3, Supplemental Tables 8 and 9)**. We then looked at lesser-known TFs predicted to be active in these lineages (**Figure 4A,B**). The top ten eRegulons in the SZC cluster at E72 shared some overlap in the target genes they putatively controlled, although they were not as highly correlated as the top 10 HC eRegulons (**Figure 4E**). *NFATC2*, for example, shares high correlation of target genes with *PEG3*, *MEF2A* and *SOX9*. The *CREB5* eRegulon also shared high similarity with the *MEF2A* eRegulon at E72 and the *DLX3* eRegulon at E59 (**Figure S5A**). eRegulons such as *NFATC2* and *MEF2A* were also active the EC1 cluster, suggesting that these may play a role in specifying epiphyseal chondrocytes to the articular chondrocyte lineage (**Figure 4B, F**). Members of the AP-1/ATF/CREB families of TFs were found at both E59 and E72 (**Figure S5C-D**). Differences between eRegulons identified at E72 versus E59 (**Figure 4, S3B, S5A-B**), could suggest temporal roles in cartilage during development.

**Figure 4.**
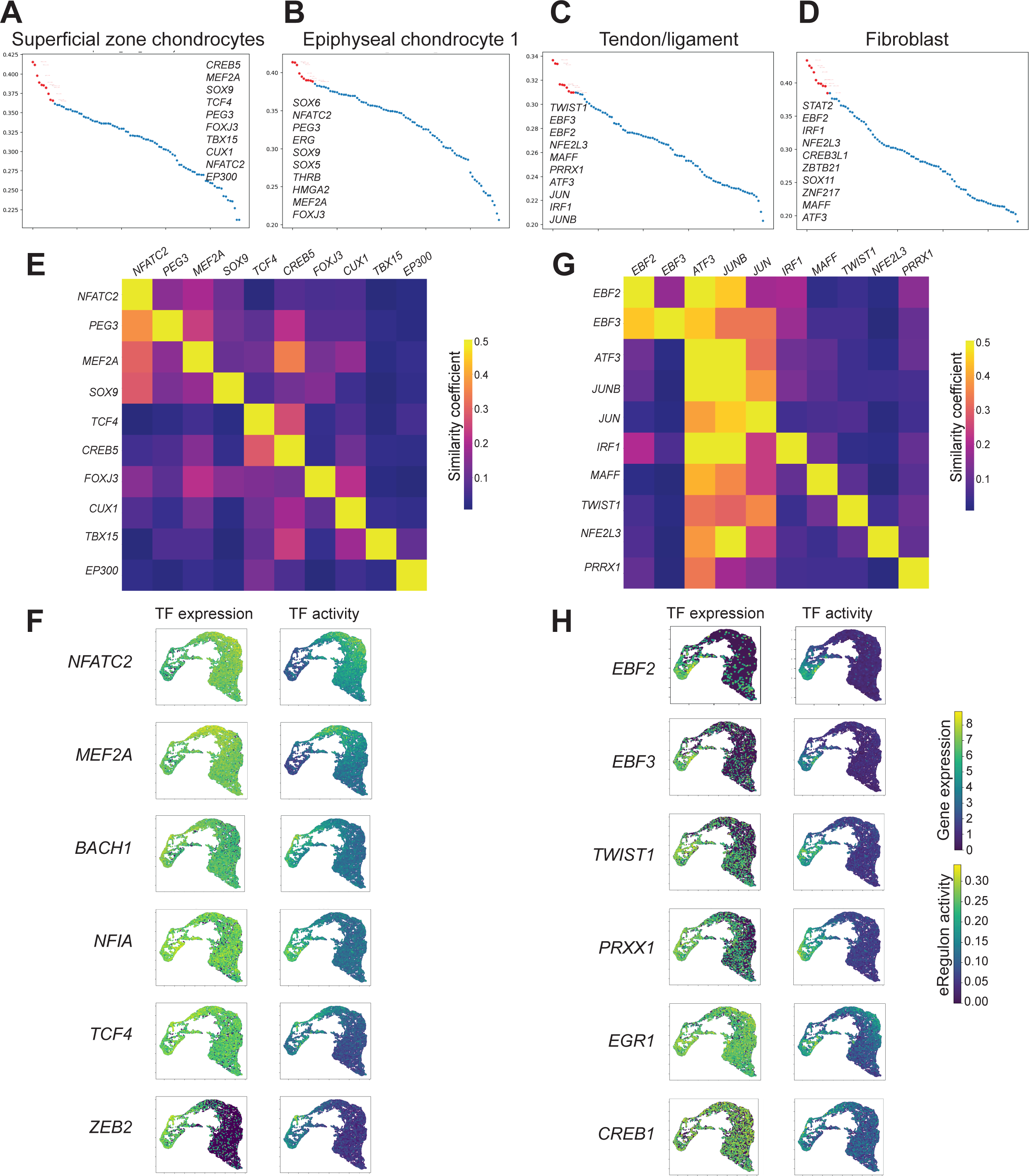
Top eRegulons in the articular cartilgae and tendon/ligament clusters at E72. (A-D) Scatter plot representing the top ten eRegulons, ranked by RSS, in SZC, EC1, Tendon/ligament, and Fibroblast clusters. (E) Correlation heatmap representing similarity between target genes putatively controlled by the top ten eRegulons in the SZC cluster. Note, differences in total target genes assigned to each TF cause heatmaps to be asymmetrical. (F) Feature plots depicting single cell-level expression (left) and activity (right) of eRegulons at E72. (G) Correlation heatmap representing similarity between target genes putatively controlled by the top ten eRegulons in the Tendon/ligament cluster. (H) Feature plots depicting single cell-level expression (left) and activity (right) of eRegulons at E72.

Our multiomic analysis of the E72 distal femur identified more eRegulons with lineage-, or cluster-based specificity, in part due to the additional cell types captured in this specimen (e.g., **Figure 4C,D,H**). EBF family members *EBF2* and *EBF3* as well as *PRRX1* and *TWIST1* were among the highest ranked eRegulons in Tendon/ligament cells, but not in chondrocytes, and were predicted to control similar target genes associated with Tendon/ligament lineages (**Tables 2 and 3, Supplemental tables 8 and 9**). Members of the AP-1 superfamily of TFs, including *JUN*, *JUNB* and *ATF3* were highly active in Tendon/ligament, Fibroblast and SZC clusters (**Figure 4C-D, S5**). It was interesting to note that several eRegulons predicted to act in SZCs, such as *DLX3*, *NFIB*, and *CREB5*, were also active in Tendon/ligament cells (**Figure 2G**).

By combining TF expression, TF motif enrichment within open chromatin regions, and the expression levels of genes associated with these regions in nuclei isolated from the distal femur at E59 and E72, we predicted eRegulons, or TF-based gene regulatory networks, governing chondrocyte subtypes and differentiation. Predicted genesets considered to be transcriptional targets of these TFs demonstrated potential cooperativity, compensatory mechanisms, and hierarchical differentiation among chondrocyte subtypes.

### Development of a model system to interrogate candidate TFs in vitro

Ideally, the importance of TFs and their eRegulons during skeletal development would be tested *in vivo* using model organisms with conditional gain and loss of function alleles. Because animals with conditional alleles may not exist for all genes of interest, we developed and validated an *in vitro* directed differentiation assay that could be used to interrogate gene function during chondrogenesis. This experimental platform was adapted from a previously published method^33,34^, and used cryopreserved chondrocytes.

Cryopreserved bi-potent human embryonic stem cell (hESC)-derived chondrocytes, isolated from 6 week old micromass tissues that were maintained in TGFβ3, retain the ability to re-generate both articular and growth plate-like cartilage tissues following thaw^34^ (**Figure 5A-B**). Replated articular and growth plate-like cartilage tissues derived from cryopreserved cells exhibited the same respective histologic morphologies as those generated with the original protocol^18,34^. To validate that differentiation of cryopreserved chondrocytes was similar to that occurring in the established protocol, we compared the transcriptomes of the original cartilage tissues to those of the replated tissues using bulk RNA-sequencing and found that the top 100 differentially expressed lineage-specific genes were similarly regulated (**Figure 5C, Supplemental Tables 12, 20**). Following this validation, we were confident that the adapted *in vitro* method using cryopreserved chondrocytes could be effectively used to study the role of TFs during human articular or growth plate chondrocyte differentiation.

**Figure 5.**
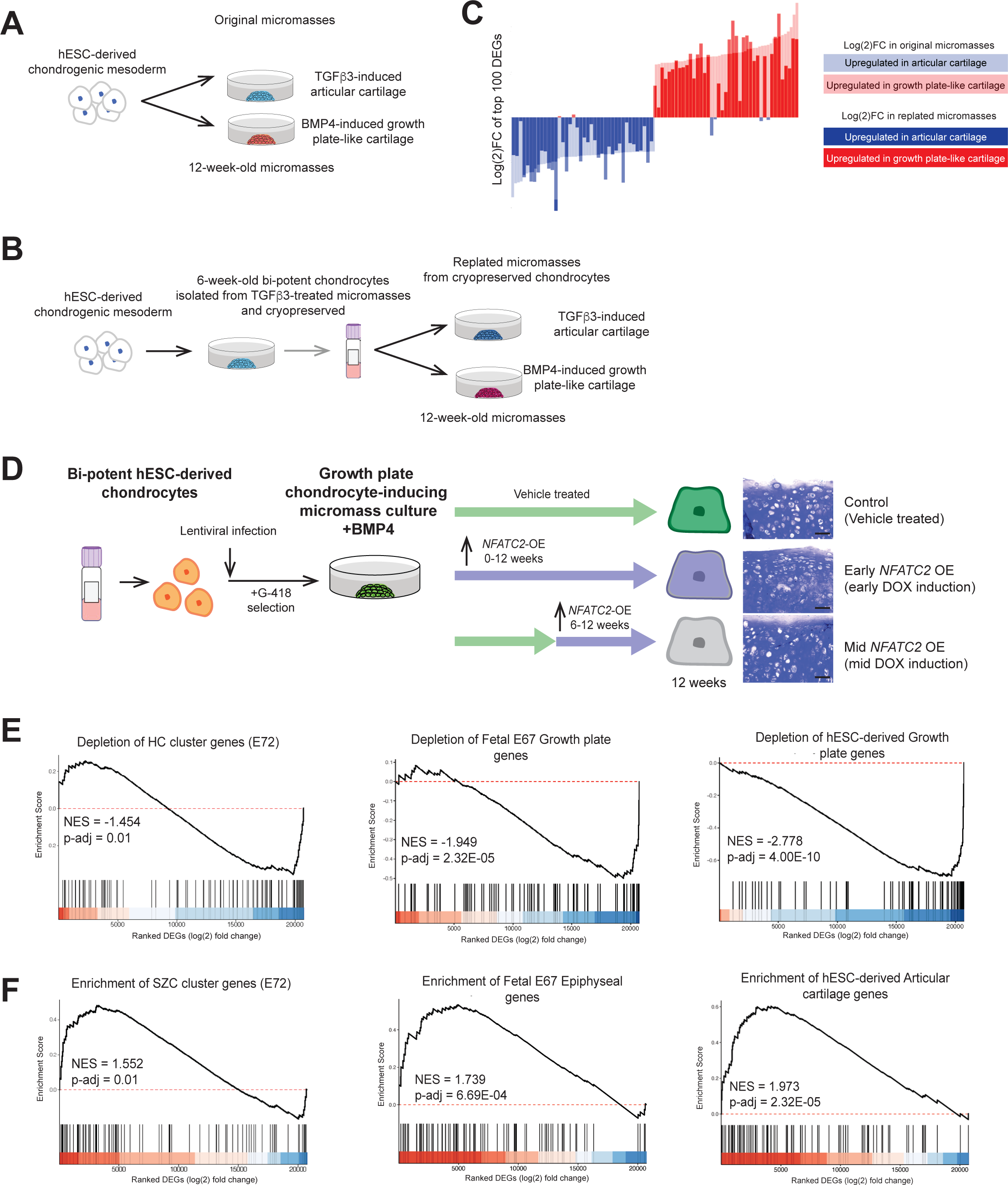
NFATC2 overexpression in in vitro growth plate cartilage results in gene expression profiles that are consistent with superficial zone and epiphyseal chondrocytes. (A) *In vitro* differentiation protocol using hESCs to generate articular and growth plate chondrocytes. (B) Modified protocol using cryo-preserved *in vitro* chondrocytes to generate articular and growth plate chondrocytes. (C) Comparison of log(2)FC values between articular and growth plate cartilage in *de novo* hESC-dervied cartilage tissues and cartilage tissues replated from cryo-preserved chondrocytes. The lighter color scheme represents comparison between *de novo in vitro* articular and growth plate chondrocytes and the darker color represents comparison between replated articular and growth plate chondrocytes. (D) Experimental design of NFATC2 overexpression, with toluidine blue staining of the resultant tissues (right). Scale bar, 100 µm (E-F) GSEA against DEGs from indicated geneset, ranked from the most up-regulated to the most down-regulated gene in early NFATC2 OE tissues compared to control vehicle-treated tissues (by log(2)FC). Genesets included (E) Top DEGs in E72 HCs, top DEGs in E67 fetal growth plate cartilage, top DEGs in hESC-derived growth plate cartilage, (F) top DEGs in E72 SZCs, top DEGs in E67 Fetal epiphyseal cartilage, and top DEGs in hESC-derived articular cartilage tissues. NES, Normalized GSEA enrichment score. p-adj, adjusted p-value using an optimized false discovery rate approach, calculated by GSEA. A positive NES indicates enrichment in the NFATC2 OE tissues compared to control tissues, and a negative value indicates depletion in the NFATC2 OE tissues.

### Overexpression of NFATC2 prevents expression of hypertrophic chondrocyte gene programs and promotes articular chondrocyte gene programs in vitro

In the E72 distal femur, *NFATC2* controls a top ranked eRegulon in SZCs and EC1 (**Figure 4**), suggesting a functional role in the specification of these cells. *In vivo*, deficiency of *Nfact2* predisposes mice to precocious articular cartilage failure^35,36^. We also found that *NFATC2* expression was significantly higher in articular chondrocytes derived from hESCs compared to growth plate-like chondrocytes^18^. We thus hypothesized that *NFATC2* promotes the formation of articular cartilage that is protected from premature degeneration. We therefore tested the effect of *NFATC2* overexpression (OE) during *in vitro* chondrogenesis using an inducible lentiviral vector (described in detail in the Methods).

*NFATC2* OE at either the early- or mid-stage of *in vitro* chondrogenesis did not affect tissue morphology (**Figure 5D**). However, RNAseq of these tissues revealed 1,571 DEGs (1,014 upregulated) after early-stage *NFATC2* OE and 732 DEGs (487 upregulated) after mid-stage *NFATC2* OE, compared to control tissues treated with vehicle alone (**Supplemental Table 13 and 14**). Gene set enrichment analysis (GSEA) showed Hallmark pathway enrichments for “Inflammatory response” and “TNFα signaling via NF-κB” in both *NFATC2* OE treatment conditions compared to control tissues (**Figure S6A,B, Supplemental Tables 15 and 16**). Assessment of Kyoto Encyclopedia of Genes and Genomes (KEGG) pathway terms also showed an enrichment for “TNF signaling pathway” and “IL-17 signaling pathway” terms in *NFATC2* OE tissues, which is consistent with known immunomodulatory roles of NFATC2 in T cells (**Figure S6C,D, Supplemental Table S17, S18**)^37,38^. Interestingly, “glycosaminoglycan biosynthesis” was depleted with early *NFATC2* overexpression, however, this did not result in differences in the presence of sulfated glycosaminoglycan (sGAG) stained metachromatically with toluidine blue in histological sections (**Figure 5D**).

Having found numerous gene expression changes caused by *NFACT2* OE, we sought to determine how this affected the expression of growth plate cartilage- and articular cartilage-associated genes. We compared *NFACT2* OE RNAseq data to the top 100 DEGs found in growth plate chondrocytes, based on data from the E72 HC and EC3 clusters in this study, and those identified in a prior study that sequenced transcriptomics of fetal growth plate cartilage and hESC-derived growth plate cartilage^18^ using GSEA (**Supplemental Table 19**). Early-stage *NFATC2* OE resulted in a significant depletion of hypertrophic chondrocyte genes from all 3 datasets (**Figure 5E** and **Figure S7A**). We next examined how OE of *NFATC2* affects the expression of genes associated with articular cartilage, and found significant enrichment of the top DEGs in the E72 SZC cluster, in fetal epiphyseal cartilage and *in vitro*-derived articular cartilage tissues^18^ (**Figure 5F**). Similar changes in gene expression patterns were observed when *NFATC2* was overexpressed in cartilage tissues that had already begun to differentiate toward the growth plate cartilage lineage (mid-stage OE; **Figure S7B-G**).

## Discussion

We leveraged multiomics to provide a snapshot of the cell types present in the distal femur at two developmental timepoints, one before and one during the development of connective tissue joint structures. In addition to identifying recognizable cell types including *PRG4*-expressing SZCs and *COL10A1*-expressing HCs, we identified epiphyseal chondrocyte subtypes that may be precursors to these more differentiated cells. EC2, found only at E59, may represent a transient cell type that has not yet committed to either lineage. At E72, we identified cell types that likely contribute to adjacent tissues, such as joint progenitor and tendon/ligament cells. This multiomic approach thus enabled us to identify differences among cells (e.g., EC1, EC2, EC3) that appear as a single cell type with either modality alone.

Using SCENIC+, which links TFs, open chromatin, and downstream target gene expression to define eRegulons, we predicted 37 active eRegulons at E59 and 94 at E72. Reassuringly, TFs for many eRegulons have recognized roles during chondrocyte differentiation; 22 were previously implicated in skeletal diseases or associated with osteoarthritis (OA) based on genome-wide association studies (GWAS). Moreover, 84 TFs are near GWAS peaks for height (**Supplemental Table 21**). Thus, other TFs-associated eRegulons identified in our study are also likely to be important for skeletal development and disease.

Our analyses also predicted novel roles of TFs that had been previously implicated in other aspects of skeletal biology. *DLX3*, along with the closely related *DLX5*, have been shown to regulate *RUNX2* during osteoblast differentiation during bone formation^39,40^. We also predicted activity of *DLX5* in HCs at E59, but the *DLX3* eRegulon was instead predicted to be highly active in SZCs at E59 and joint progenitors at E72. Additionally, target genes of *DLX3* show high correlation with those targeted by *CREB5* at both timepoints, suggesting that these TFs work cooperatively to control gene programs in the superficial zone of developing human cartilage. In a similar vein, *MEF2A* was predicted to be active in the SZCs in the E72 distal femur, in contrast to its reported positive regulation of the HC-associated gene *COL10A1*, encoding type X collagen^41^. While it is possible that *DLX3* and *MEF2A* control different sets of genes in HCs that were not captured by the SCENIC+ algorithm, these data lead us to hypothesize a novel role for these TFs in SZCs and joint cell specification during human chondrogenesis. Previous studies did not report articular cartilage abnormalities in knockout mice^42,43^; however, severe non-skeletal phenotypes may have limited detailed studies of the cartilage.

SCENIC+ enabled us to predict target genes of different TFs, and whether these were shared among two or more TFs. A high similarity of target genes could indicate co-operativity between TFs. The open chromatin regions, or putative enhancers, near these target genes should be evaluated in functional reporter assays, to determine whether each TF binding motif is required or sufficient for expression of the target gene. Shared transcriptional targets could also indicate redundancy, and downstream knockout and rescue experiments can help determine the necessity of these TFs in chondrogenesis and lineage-specificity. eRegulons in HCs and PHCs were predicted to have high similarity in their target genes. This could reflect the coordination of terminal differentiation of the HCs in the growth plate. The chondrocytes of the future articular cartilage (e.g., SZCs, EC1), in contrast, have not yet been specified to their end-stage cell types. These epiphyseal chondrocytes likely have other potential fates, including permanent articular chondrocytes lining the joint surface, or growth plate-like chondrocytes that participate in the formation of the secondary ossification center. The dynamic nature of these cell types therefore might require that master TFs coordinate different genesets.

Our approach uncovered eRegulons for emerging tendon and ligament cells captured at E72, including CREB and EBF family members. Several SZC eRegulons were also detected in tendon/ligament cells, such as *CREB5*. CREB5 may therefore be more intimately involved in joint tissue development, perhaps through the regulation of synovial, ligament or tendon fibroblast populations^44,45^. Our independent discovery of conserved EBF and CRE binding motifs in functional putative enhancers of *scxa/Scx/SCX* in zebrafish, mouse and human suggests that these multiomic data have the potential to uncover novel gene regulatory networks upstream of this earliest known tenocyte marker (Niu et al., *Dev Cell, in press*). The overlap in transcriptional activity between SZC and tendon/ligament cell types could also be attributed to a common progenitor within the developing joint^46^. Other TFs, including the well-known limb bud gene *PRRX1*, represent hitherto unknown networks controlling tendon specification.

Because eRegulons are computationally predicted, their functional relevance must be tested. We developed an experimental platform that uses cryo-preserved chondrocytes derived from human pluripotent stem cells, which is comparable to but faster than beginning with undifferentiated stem cells. Using this platform, we demonstrated that *NFATC2* OE favors articular chondrocyte differentiation, which is consistent with the reduced stability of articular cartilage and increased expression of the HC gene *COL10A1* in *Nfatc2* knockout mice^35,36,47,48^. These results suggest that NFATC2 may play an active role in lineage commitment of chondrocytes toward epiphyseal/articular chondrocyte fates during development *in vivo* and may repress the transdifferentiation of articular chondrocytes to HCs postnatally, as observed in OA pathology. Thus, we demonstrated the utility of this *in vitro* model for testing the functional roles of other genes and eRegulons during human chondrogenesis such as those uncovered herein.

We appreciate that single nucleus multiomic methods lack spatial resolution, which will be needed to confirm predicted relationships between the EC1 and SZC cluster, the EC3 with PHC cluster, and articular chondrocyte and tendon/ligament cell clusters. We were also limited by the availability of donor tissue to studying only 2 developmental timepoints from single specimens. Hence, some predicted eRegulons could be false positive findings that would only be recognized by adding more samples or by functionally testing their roles *in vivo*. Despite these limitations, single cell resolution of the transcriptomic and epigenetic profiles of the cells that comprise the distal femur enabled us to predict master regulators of chondrogenic and joint cell differentiation. These *in vivo* data, combined with the *in vitro* experimental platform we established, will facilitate rapid prioritization of genes and pathways that should be further studied for temporal and cell-type specific roles in human developmental chondrogenesis.

## Methods

### Dissection of fetal samples

Human fetal donor samples were collected with consent from first trimester terminations on embryonic day (E)59 and E72 at the University of Washington (UW) Birth Defects Research Laboratory (BDRL) in full compliance with NIH ethical guidelines, and with the approval from the UW Review Boards for the collection and distribution of human tissues for research, and from Harvard University and Boston Children’s Hospital for the receipt and use of such materials. The samples were briefly washed in Hank’s Balanced Salt Solution (HBBS) and transported in the same buffer at 4°C during overnight shipment. Cartilaginous tissue from the distal femur was dissected under a light dissection microscope in 1x phosphate buffered saline (PBS) on ice, and soft tissue was removed. The samples were micro-dissected to enrich for chondrocytes, excluding cells from the distal ossification front of the femur, but including a subset of connective tissue cells, which were then isolated enzymatically for further study. Knee joints from the contralateral limb (E72) or a stage matched fetal donor (E59) were fixed in 10% formalin and processed for paraffin embedding.

### 10x Multiome Library Preparation

Distal femur tissues were enzymatically digested in 0.2% type I collagenase (Corning) for up to 4 hours at 37°C. The cells were pelleted by centrifugation and the supernatant was removed. Cells were lysed and nuclei were isolated following manufacturers’ instructions as detailed in Nuclei Isolation for Single Cell Multiomics ATAC + Gene Expression Sequencing protocol (CG000365-Rev C), with lysis time varying based on developmental stage of the sample (5 minutes for E59, 7 minutes for E72). Single nuclei ATAC and RNA sequencing libraries were prepared using a Chromium Single Cell Multiome ATAC + Gene Expression kit, which included reagents and protocols found in PN-1000285.

In brief, isolated nuclei were transposed and the resultant suspension was loaded into a microfluidics chip and run on a Chromium controller to generate Gel Beads-in-emulsion (GEM) containing single nuclei. Incubation of GEMs resulted in 10x barcoded DNA from transposed fragments and 10x barcoded cDNA from RNA. Single cell ATAC and RNA seq libraries were prepared following manufacturers’ instructions. RNA and ATAC-seq libraries pooled together and sequenced. The E59 distal femur sample was sequenced on an Illumina HiSeq and the E72 distal femur sample was sequenced on an Illumina NovaSeq at the Bauer Core Facility at Harvard University.

### Single cell RNA and ATAC sequencing data processing and analysis Mapping and quality control

Raw sequencing data were processed using Cellranger (v2.0.0). Raw sequencing reads were mapped to the GRCh38 human genome build and subsequently analyzed using Seurat v4.3^21^ and Signac v1.10^49^. The resultant data were filtered based on number of unique genes per cell (>1000), distribution of RNA counts (log10 genes per unique molecular identifier > 0.7), mitochondrial ratio (< 20%), nucleosome signal (< 2), and TSS enrichment (>1).

### RNA sequencing analysis

Cell cycle phases were scored (CellCycleScoring). RNA data were normalized and scaled using the SCTransform function, controlling for mitochondrial content, ribosomal content, and cell cycle phase. PCA was performed based on variable features. UMAP using the first 30 principal components was performed and cells were clustered using a smart local moving (SLM) algorithm at resolution 0.8. Plots were generated using DimPlot, FeaturePlot, DotPlot, and DoHeatmap functions.

### ATAC sequencing analysis

The accessible chromatin regions were identified using MACS2^50^ (v2.1.1). The data was then normalized, and dimensionality reduction was conducted using latent semantic indexing (FindTopFeatures, RunTFIDF, RunSVD). UMAP was performed using 2-50 LSI components. Differentially accessible peaks between named clusters were calculated using the FindAllMarkers function using a logistic regression test, returning only positively expressed markers with a log2 fold change > 0.1 and Bonferroni-adjusted p value <0.05. Plots were generated using the CoveragePlot function.

### Integrated RNA and ATAC-seq analysis

Integrated analysis of RNA and ATAC seq data was conducted using weighted nearest neighbors (wKNN), using the first 50 PCs and 2-50 LSI components. UMAP was performed and cells were clustered using the SLM algorithm at resolution 0.8. Clusters were then named based on DEGs and expression levels of known markers. For the E72 sample, off target cell populations including neuronal, hematopoietic, and skeletal muscle cells were filtered out and the analysis was re-run. Differentially expressed genes and differentially accessible peaks between named clusters were calculated using the FindAllMarkers function using a logistic regression test, returning only positively expressed markers with a log2 fold change > 0.1 and Bonferroni-adjusted p value <0.05.

### Transcription factor activity analysis

Single cell TF motif activity was estimated using chromVAR for a set of 760 TFs from JASPAR’s 2020 CORE vertebrate collection^28,51^. Differential motif activity was calculated using the FindAllMarkers function (log2 fold change >0.1 and Bonferroni-adjusted p value <0.05).

### Identification of putative regulatory networks by SCENIC+ Identification of eRegulons

The SCENIC+ algorithm consists of three major tasks: identification of enhancer regions with shared accessibility patterns, prediction of TF binding sites, and generation of eGRNs by combining TF expression, TF motif enrichment, target region accessibility, and target gene expression. The computational steps are briefly described below.

Consensus peak calling was performed using pycisTopic. Iterative peak calling was performed using MACS2 to generate a obtain a consensus peak set for each cell type separately and then for the union of peaks across all cell types. Topic modeling and identification of DARs was then performed as described in Gonzalez-Blas et al., 2023^32^. TF binding sites were then predicted using pycisTarget, using the previously generated cisTarget database (https://resources.aertslab.org/cistarget/). TF-to-gene importance scores were calculated by predicting TF expression using raw gene expression. Region-to-gene importance scores were calculated by predicting TF expression from region accessibility, for all regions around a gene’s search space. The gene search space was defined as a minimum of 1kb and maximum of 150kb both upstream and downstream of the nearest gene or promoter, and the promoter of a gene was defined as the TSS +/- 10 bp, the default parameters used in SCENIC+. All regions enriched for a TF motif and all genes linked to these regions were used to generate TF-region-gene triplets. Target genes of eRegulons were calculated by gene set enrichment analysis (GSEA) as described in Gonzalez-Blas et al., 2023^32^, and eRegulons with fewer than 10 target genes or with target genes obtained from negative region-gene correlations were discarded. The list of all putative eRegulons, along with their predicted targets, can be found in **Supplemental Tables S8** and **S9.**

eRegulon enrichment was then calculated based on area under the curve (AUC) of ranked genes and consensus peaks, where target genes or peaks were considered enriched at AUC at 5% of ranking. The eRegulon enrichment score for genes were normalized and used to perform UMAP, using Python’s UMAP v.0.5.2. The resultant plots for UMAP visualization and eRegulon enrichment were generated using matplotlib. eRegulon specificity scores were calculated for each cell type and each eRegulon using the RSS algorithm described in Gonzalez-Blas et al., 2023. The RSS scatter plot was generated using matplotlib. The correlation between genes or regions controlled by eRegulons was calculated using the normalized intersection metric and the resultant heatmap plotted using seaborn.

### In vitro functional studies using chondrocytes derived from human pluripotent stem cells Generation of chondrocytes from hESCs

All reported research involving human embryonic stem cells was approved by IRB (IRB-P00017303) and ESCRO (ESCRO-2015.4.24) regulatory bodies at Boston Children’s Hospital. Detailed descriptions of this directed differentiation protocol have been published^18,33,34^. In brief, embryoid bodies (EBs) were generated from H9 hESCs (Wicell) and cultured in suspension in the presence of BMP4 (1 ng/mL) and Y-27632 (5 μM) in Stem-Pro-34 media (Life Technologies) supplemented with L-glutamine, L-ascorbic acid, monothioglycerol, and transferrin. On day 1, EBs were collected and resuspended in Stem-Pro media with Activin A (2 ng/mL), bFGF (5 ng/mL), BMP4 (3 ng/mL), and Y-27632 (5 µM) to induce primitive streak-like mesoderm. After 44 hours, the EBs were harvested, cells were dissociated using TrypLE (Life Technologies) and cultured as monolayers in 96-well tissue culture plates in Stem-Pro media containing bFGF (20 ng/mL), SB431542 (5.4 µM), dorsomorphin (4 µM), and IWP2 (2 µM). After 48 hours, on day 5, monolayer cultures were maintained in Stem-Pro media containing bFGF (20 ng/mL) until day 14. Cells were maintained in hypoxic conditions (5% O_2_/5% CO_2_/90% N_2_) for 11 days and 5% CO_2_/atmospheric O_2_ conditions for the remaining period. Cartilage tissues were generated from monolayer cultures on day 14 by plating cells in micromass culture. For each micromass, 250,000 cells were seeded on Matrigel-coated 24-well plates in base chondrogenic media consisting of high glucose DMEM supplemented with 1% ITS, L-proline (40 μg/mL), Pen/Strep, dexamethasone (0.1 μM), L-ascorbic acid (100 μg/mL) and TGFβ3 (10 ng/mL) for 2 weeks. This protocol was used to generate articular cartilage (by maintaining cells in TGFβ3 for 10 additional weeks) or growth-plate like cartilage (by maintaining cells in base chondrogenic medium supplemented with BMP4 (50 ng/mL)).

Chondrocytes were isolated from 6-week-old tissues maintained in TGFβ3 using 0.2% type I collagenase for 2 hours at 37°C. Isolated chondrocytes were cryopreserved in media consisting of 50% heat-inactivated fetal bovine serum (FBS, Corning), 40% Iscove’s Modified Dulbecco’s Medium (IMDM), and 10% dimethyl sulfoxide and stored in liquid nitrogen until further use.

### Generation of lentiviral doxycycline-inducible NFATC2-overexpression vector

The 2776 bp long *NFATC2* coding sequence (CDS) was PCR-amplified from cDNA preparations from 12-week-old hESC-derived articular cartilage using the Platinum SuperFi II PCR Mastermix (Gibco) (Forward primer: CACCATGAACGCCCCCGAGC, Reverse primer: TTACGTCTGATTTCTGGCAGGAGGTCC) and cloned into a Gateway vector (pENTR-*NFATC2*) using the pENTR/SD/D-TOPO cloning kit (Gibco). NFATC2 CDS was transferred into the pInducer20 lentivirus destination vector (Addgene Plasmid #44012^52^) using Gateway LR Clonase to generate pInducer20-*NFATC2*.

### Production of lentivirus

HEK293T cells were maintained in 10 cm tissue culture-treated plates in DMEM/F12 supplemented with 10% heat inactivated FBS at 37°C until ∼70% confluent. Lentiviral expression plasmid (pInducer20-*NFATC2*) (6 µg), psPAX2 (4.5 µg), and pMD2.G (1.5 µg) plasmids were added to a tube containing 500 µL Opti-MEM (Gibco). Diluted Fugene (36 µL of Fugene in 500 µL Opti-MEM) was added dropwise to the tube containing the plasmid solution. The mixture was allowed to incubate for 15 minutes at room temperature to form the DNA-Fugene complex. The complex was then diluted in OptiMEM and 6 mL was added to every 10 cm tissue culture plate with HEK293T cells. One day later, the media was replaced with base chondrogenic media (described above). Supernatant containing virus was collected in sterile tubes at 48 and 72 hours post transfection. Cell debris was removed with centrifugation at low speed and filtration through 0.45 µm PES membrane filters. Supernatant containing virus was stored at -80°C until further use.

### Generation of cartilage tissues with overexpression vector

Cryopreserved chondrocytes derived from hESCs were thawed and plated in monolayer culture on 24-well gelatin-coated tissue culture-treated plates at 250,000 cells/well. Monolayer cultures were maintained in base chondrogenic media supplemented with L-ascorbic acid (100 µg/mL) and heat-inactivated FBS (2% of media) and allowed to expand over three days. On day 4, monolayers were transduced with lentivirus (500 µL of virus-containing supernatant). After 24 hours, monolayers were washed twice with IMDM, and cultured in base chondrogenic media supplemented with L-ascorbic acid (100 µg/mL) and 2% FBS. After 24 hours, infected monolayers underwent selection by treatment with Geneticin (G-418; 500 μg/mL; Gibco) until all uninfected control cells were dead.

Lentiviral-infected chondrocytes were dissociated using 0.25% Trypsin-EDTA and 250,000 cells were plated in micromass (see above) in maintained in base chondrogenic media supplemented with L-ascorbic acid (100 µg/mL) and TGFβ3 (10 ng/mL) for 2 weeks. To induce growth plate chondrocyte differentiation, TGFβ3 was replaced with BMP4 (50 ng/ml) in the media after two weeks of micromass culture. For early *NFATC2*-overexpression, doxycycline (1 µg/ml) was administered to micromass cultures starting at 3 days after seeding. For mid-stage *NFATC2*-overexpression, doxycycline was administered starting at 6 weeks of micromass culture. Chondrogenic media containing BMP4 and doxycycline, if indicated, was replaced every 3-4 days. Micromass tissues were collected after 12 weeks of culture for transcriptomic and histologic analyses.

### Bulk RNA-sequencing

Cartilage tissues were enzymatically digested with 0.2% type I collagenase for up to 2 hr at 37°C to release single cells. Total RNA from isolated cells was purified using silica column-based kits. RNA quality and quantity were assessed via Bioanalyzer (Agilent), and samples with RIN values greater than 7 were used in subsequent library preparations (See Metadata in **Supplemental Table 20**). RNA sequencing libraries were prepared using the NEBNext Ultra II RNA Library Prep kit (Azenta) and sequenced (Illumina) to a minimum of 30,000 reads per sample using 2 x 150bp paired-end reads.

### Bulk RNA-seq processing and analysis Mapping, normalization, and differential testing

Paired-end sequencing reads were mapped to the GRCh38 human genome build using Salmon v1.8.0 with the following parameters in the quant function (-l A,--validateMappings, --gcBias, --numBootstraps 100). Salmon quantification files were imported to R v4.2.1 using tximport library v1.24.0. Transcript counts were summarized at the gene level using the GTF file mapping. Counts data was loaded into DESeq2 (v. 1.38) and samples were batch-corrected using Combat-Seq. Differential gene testing was conducted using the DESeq function (alpha = 0.05) and p-adjusted values were calculated using the Benjamin-Hochberg FDR method. DEGs were defined as those genes having a p-adj value < 0.05.

### Heatmap generation

To visualize expression patterns, curated lists of select genes were subset from normalized counts and visualized using the base ‘heatmap’ function (scaled by row). Significance for differences in gene expression was set at p-adj value < 0.05.

### Gene-set enrichment analysis

DEGs were tested for enrichment in Kyoto Encyclopedia for Genes and Genomes^53^ (gseaKEGG), and Molecular Sigantures Database (MSigDB) Hallmark pathways^54^ (gseaHallmark) annotated terms using clusterProfiler (v4.2.0). All genes in the reference transcriptome were used for GSEA, following quality control. Only terms with at least 10 genes in the set with a p-adjusted value cutoff of 0.05 were extracted.

### GSEA of lineage-specific genesets

Genesets representing SZC, EC1, EC3 and HCs were generated using the top 100 DEGs by Log(2)FC in each respective E72 cluster. The top 100 DEGs in each lineage from the published bulk RNA sequencing comparison of E67 fetal epiphyseal cartilage and growth plate cartilage, and of hESC-derived articular versus growth plate-like cartilage^18^ were also used as lineage-specific genesets for comparison. GSEA was performed with ranked (by Log(2)FC) gene expression in early-stage *NFATC2* OE or mid-stage *NFATC2* OE tissues compared to control tissues using GSEA v4.3.3^55^.

### Shared directional gene expression

To compare the original in vitro derived micromass tissues to those that were replated from cryopreserved bi-potent chondrocytes, the top 100 DEGs (50 from each lineage) from the original hESC-derived growth plate and articular cartilage transcriptomic comparison were extracted and the log(2)FC values in both the original micromasses (**Supplemental Table 17**) and the cryopreserved and replated micromasses (**Supplemental Table 10**) were plotted in bar plots using *ggplot2*.

### Histology

Fetal E59 and E72 knee joints and *in vitro*-derived cartilage tissues were fixed in 10% formalin and embedded in paraffin using standard procedures. 5 µm sections were stained with toluidine blue to visualize sulfated glycosaminoglycans.

## Data Accessibility Statement

Data accessibility statement

The raw snRNA-seq, snATAC-seq data and bulk RNA-seq data will be available on GEO upon publication of the peer-reviewed manuscript. Raw data for bulk RNA sequencing samples used as reference genesets were previously published^18^ and can be found in GEO accession GSE195688.

## Acknowledgements

We would like to thank the University of Washington Birth Defects Research Laboratories, supported by NIH award number 5R24HD000836 from the Eunice Kennedy Shriver National Institute of Child Health and Human Development; the funding sources for this work (NIAMS - R01-AR081274 (AMC and TDC), R01-AR073821 (AMC); the Harvard Bauer Core Facility for NGS support; pInducer20 was a gift from Stephen Elledge (Addgene plasmid #44012); the Bone cell RNA-seq and Directed differentiation cores of the Center for Skeletal Research NIH-funded program (P30-AR075042), and Dr. Matthew Warman, whose constructive comments helped to improve this manuscript.

## Author Contributions

All authors agree to attest to the accuracy and integrity of the work and approved the final version of the manuscript for submission. Conceptualization (AMC and TDC); data curation and resources (DV, GS); formal analysis, investigation, and methodology (DV); funding acquisition (AMC and TDC), supervision (AMC), writing–original draft preparation (DV and AMC), writing–review and editing (DV, GS, TDC, AMC).

## Figure Legends

**Figure 3. Cell cluster relationships based on the activity of eRegulons (TFs) were identified using enhancer-based gene regulatory network analysis, SCENIC+.**

(A-B) UMAP projection of cells and cell cluster assignments based on eRegulon activity at E59 (A) and E72 (B). Note, the wKNN cell cluster names projected onto this eRegulon UMAP are shown in Supplemental Figure 4A-B.

(C-E) Feature plots depicting single cell-level expression (left) and activity (right) of known transcription factors/eRegulons identified in both timepoints at E59.

(F-H) Feature plots depicting single cell-level expression (left column) and activity (right) of known, shared transcription factors/eRegulons at E72. These include articular chondrocyte TFs CREB5 and ERG (C,F); chondrogenic TFs SOX5, SOX6, and SOX9 (D,G); hypertrophic chondrocyte TFs RUNX2 and MEF2C (E,H). Arrows point to cell clusters with eRegulon activity, unless activity is detected in all cells (no arrows shown).

## Supplemental Figure Legends

**Supplemental Figure 1.**
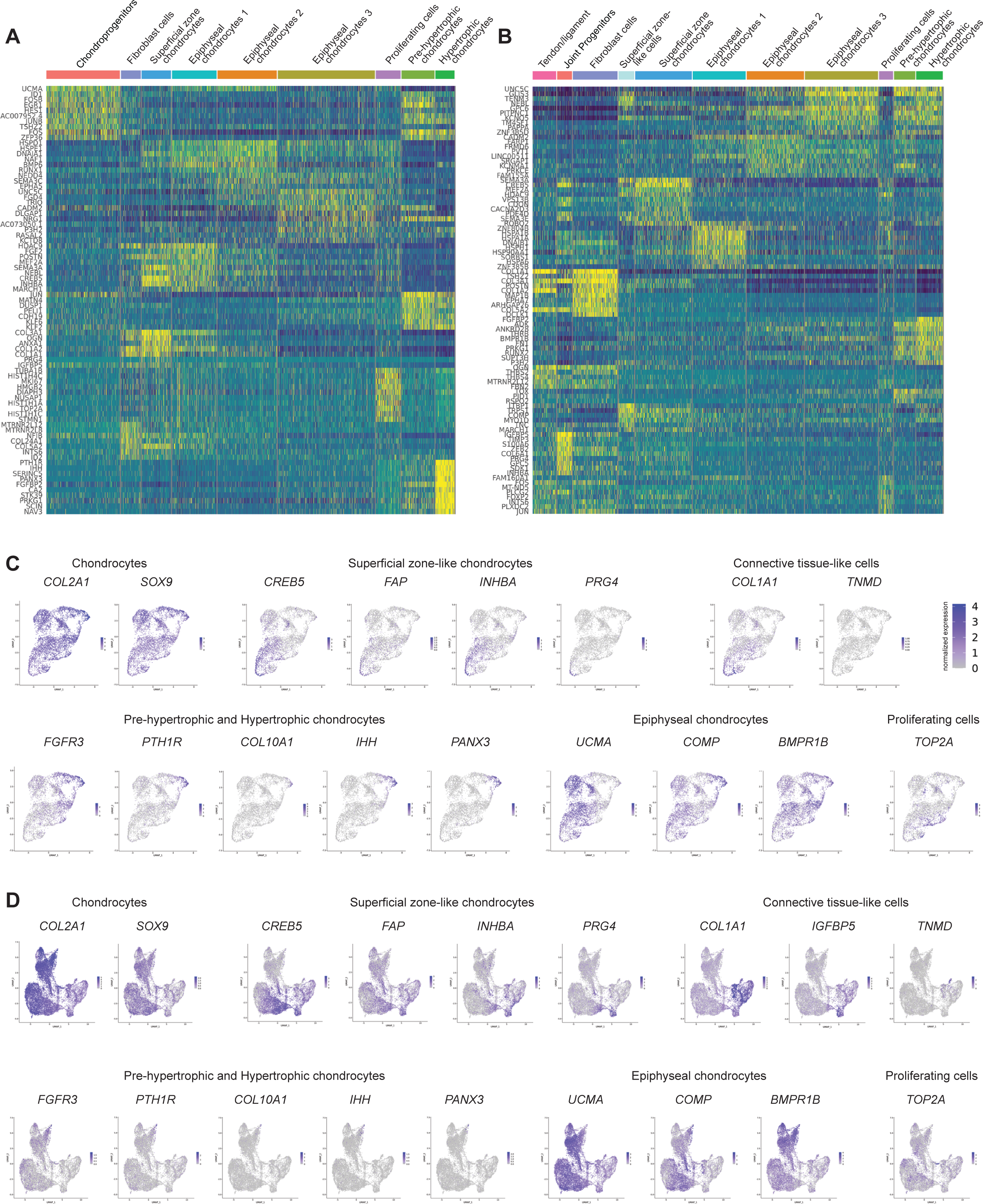
Gene expression in cell clusters. (A-B) Heatmaps represent expression of top 10 DEGs by Log(2)FC in each cell cluster at E59 (A) and E72 (B). (C-D) Feature plots depict normalized gene expression of select marker genes in cells at E59 (C) and E72 (D).

**Supplemental Figure 2.**
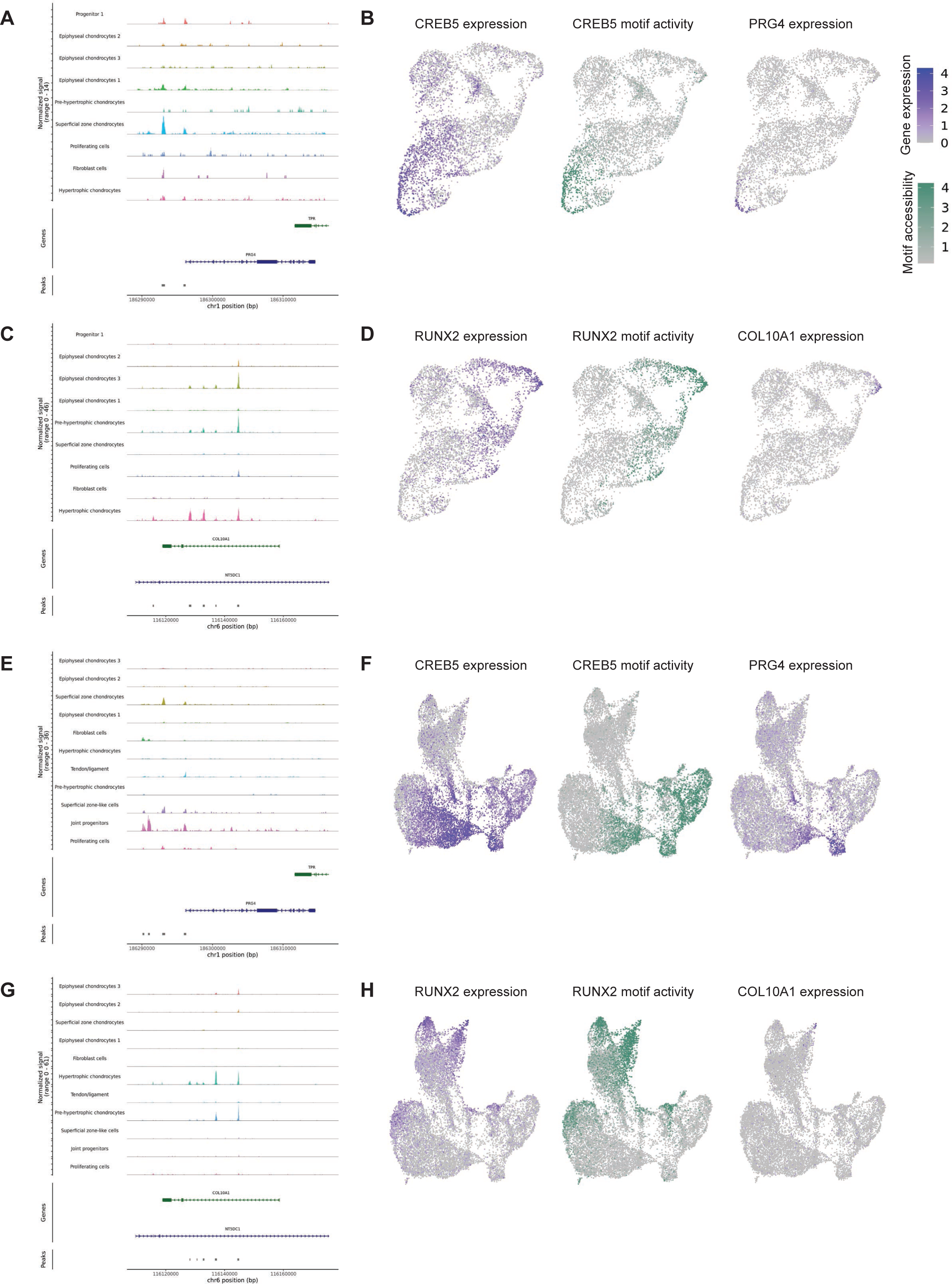
Representative open chromatin regions, motif accessibility and target gene expression. (A) Genome browser tracks around the PRG4 locus at E59. (B) Features plots for CREB5 expression (left), CREB5 motif accessibility (middle), and PRG4 expression (right) at E59. (C) Genome browser tracks around the COL10A1 locus at E59. (D) Features plots for RUNX2 expression (left), RUNX2 motif accessibility (middle), and COL10A1 expression (right) at E59. (E) Genome browser tracks around the PRG4 locus at E72. (F) Features plots for CREB5 expression (left), CREB5 motif accessibility (middle), and PRG4 expression (right) at E72. (G) Genome browser tracks around the COL10A1 locus at E72. (H) Features plots for RUNX2 expression (left), RUNX2 motif accessibility (middle), and COL10A1 expression (right) at E72.

**Supplemental Figure 3.**
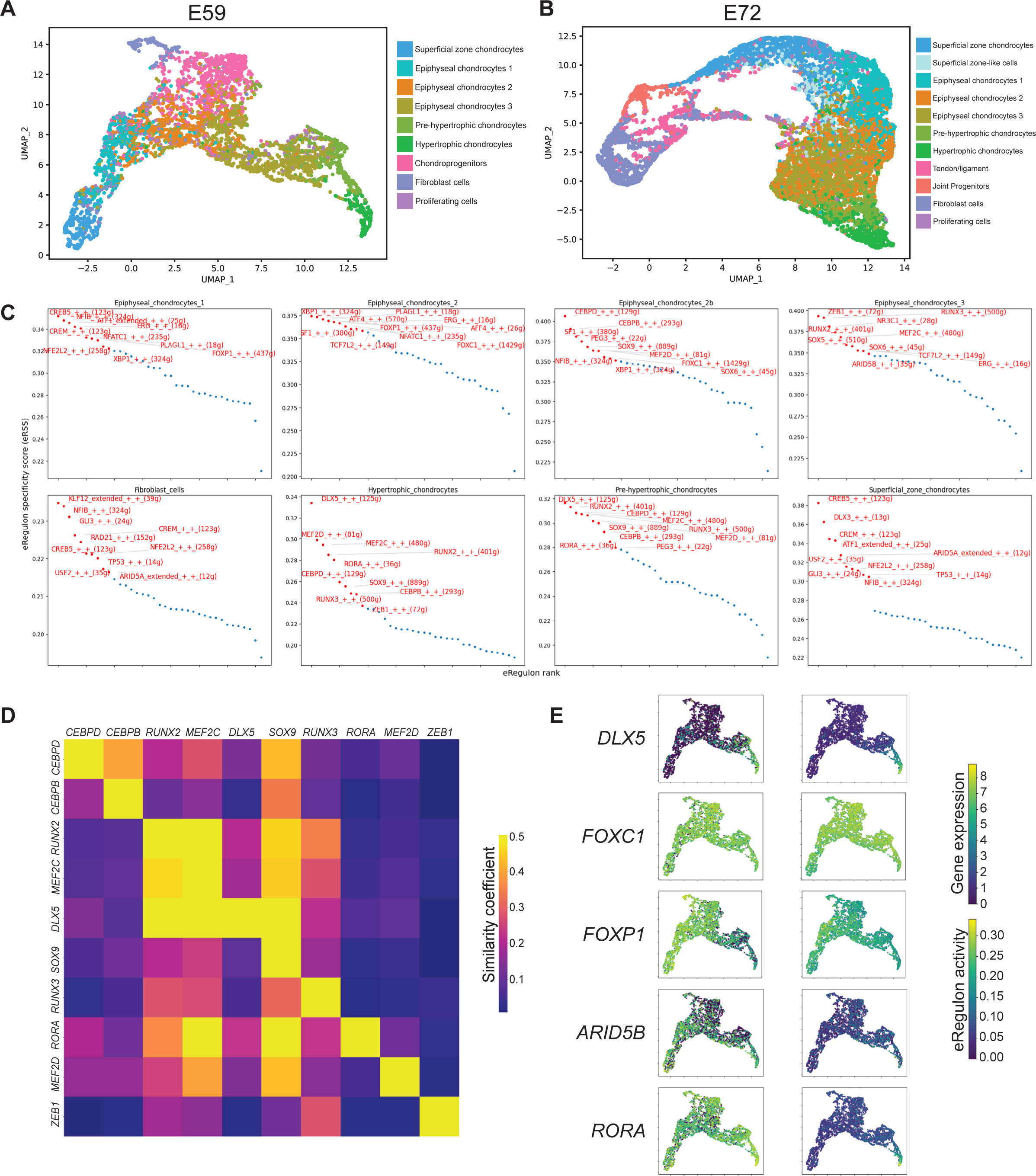
Distribution of previously identified cell types in eRegulon-based UMAP and the top ranked eRegulons based on specificity of activity in each eRegulon-defined cell cluster at E59. (A) eRegulon-based dimensionality reduction UMAP with E59 cells colored/labeled by their original WNN identity. (B) eRegulon-based dimensionality reduction UMAP with E72 cells colored/labeled by their original WNN identity. (C) Top ten eRegulons/TFs, ranked by their eRegulon specificity score (RSS), in each cell type cluster at E59. ERegulon specificity scores are on the y-axis and the numbered rank of each TF/eRegulon is on the x-axis. (D) Correlation heatmap representing similarity between target genes putatively controlled by the top ten eRegulons in the HC cluster. Note, differences in total target genes assigned to each TF cause heatmap to be asymmetrical. (E) Feature plots depicting single cell-level expression (left) and activity (right) of eRegulons.

**Supplemental Figure 4.**
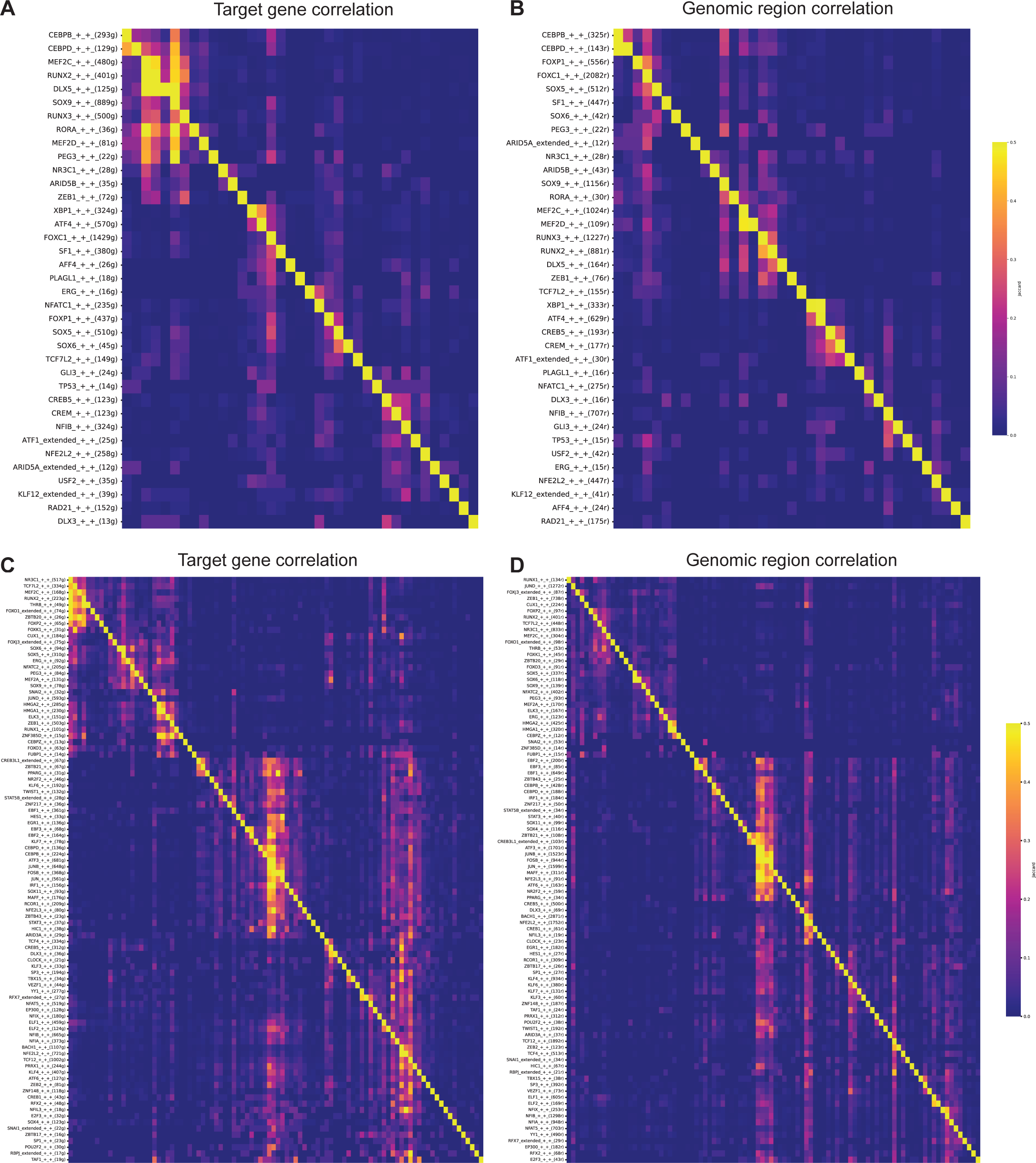
Correlation of genes and regulatory regions controlled by eRegulons. (A-B) Correlation of genes (A) and regions (B) controlled by eRegulons in the E59 distal femur. (C-D) Correlation of genes (C) and regions (D) controlled by eRegulons in the E72 distal femur

**Supplemental Figure 5.**
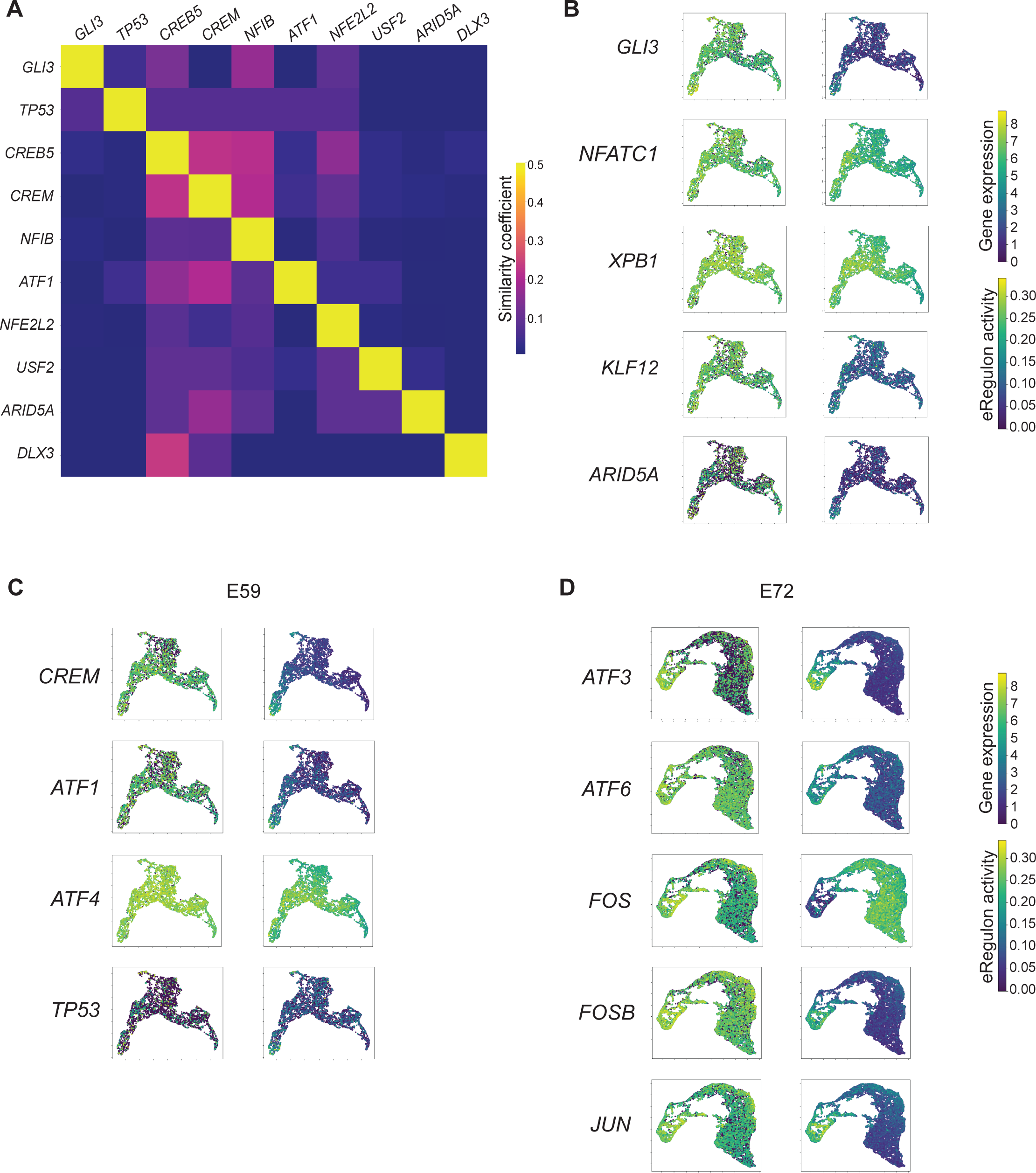
Illustration of eRegulon activity in the articular cartilage lineage at E59 and E72. (A) Correlation heatmap representing similarity between target genes putatively controlled by the top ten eRegulons in the SZC cluster at E59. Note, differences in total target genes assigned to each TF cause heatmap to be asymmetrical. (B) Feature plots depicting single cell-level expression (left) and activity (right) of eRegulons at E59. (C,D) Feature plots depicting single cell-level expression (left) and activity (right) of representative eRegulons in the AP-1/CREB/ATF transcription factor families at E59 (C) and E72 (D).

**Supplemental Figure 6.**
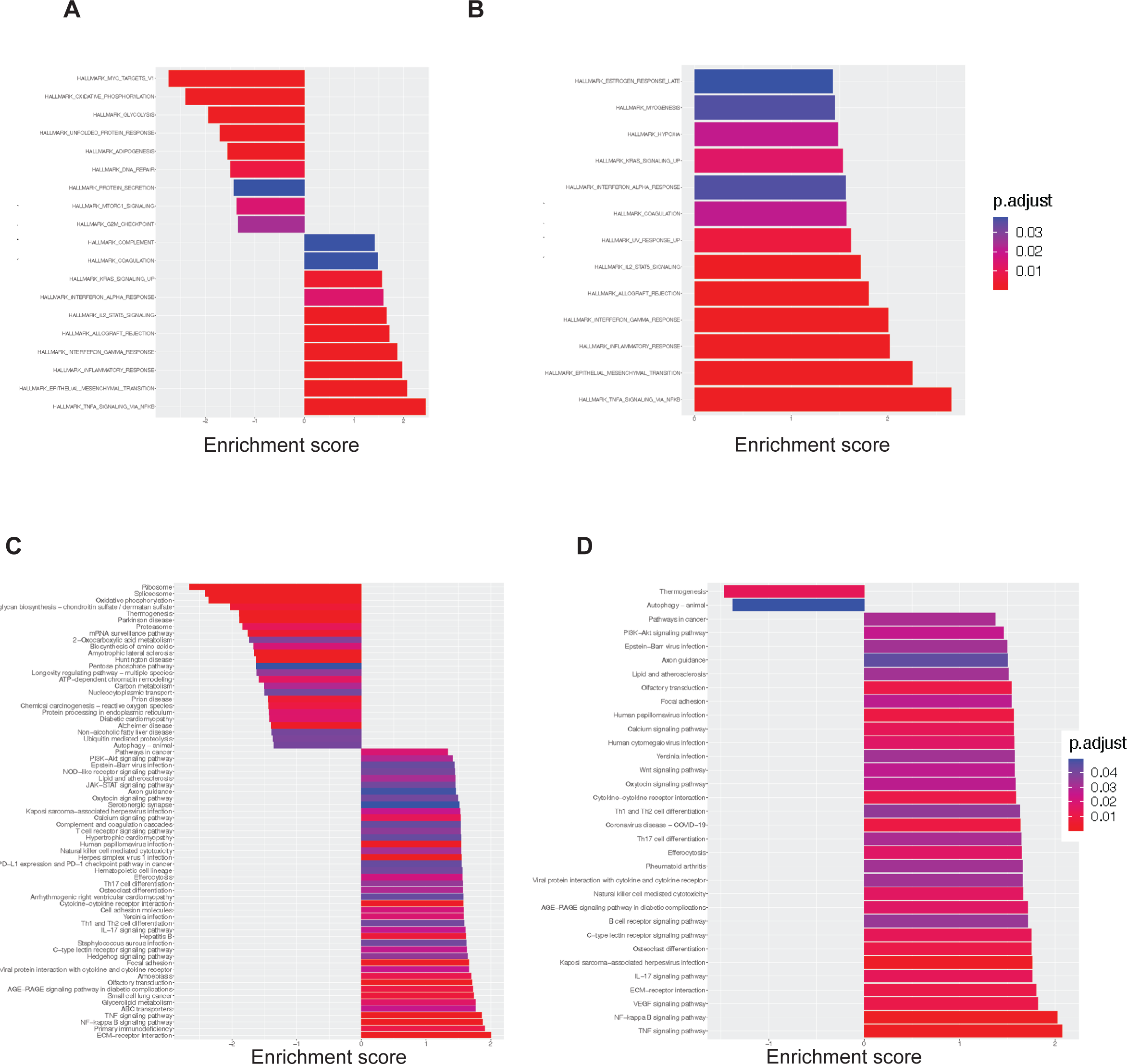
Gene set enrichment analysis of NFATC2 OE versus control tissues. (A,B) Gene set enrichment analysis against Hallmark pathway terms in the early NFATC2 OE (A) and mid NFATC2 OE (B) samples compared to control. (C,D) GSEA against KEGG terms in the early NFATC2 OE (C) and mid NFATC2 OE (D) samples compared to control. The color value represents p-adjusted value. A positive enrichment score indicates upregulation in NFATC2 OE samples.

**Supplemental Figure 7.**
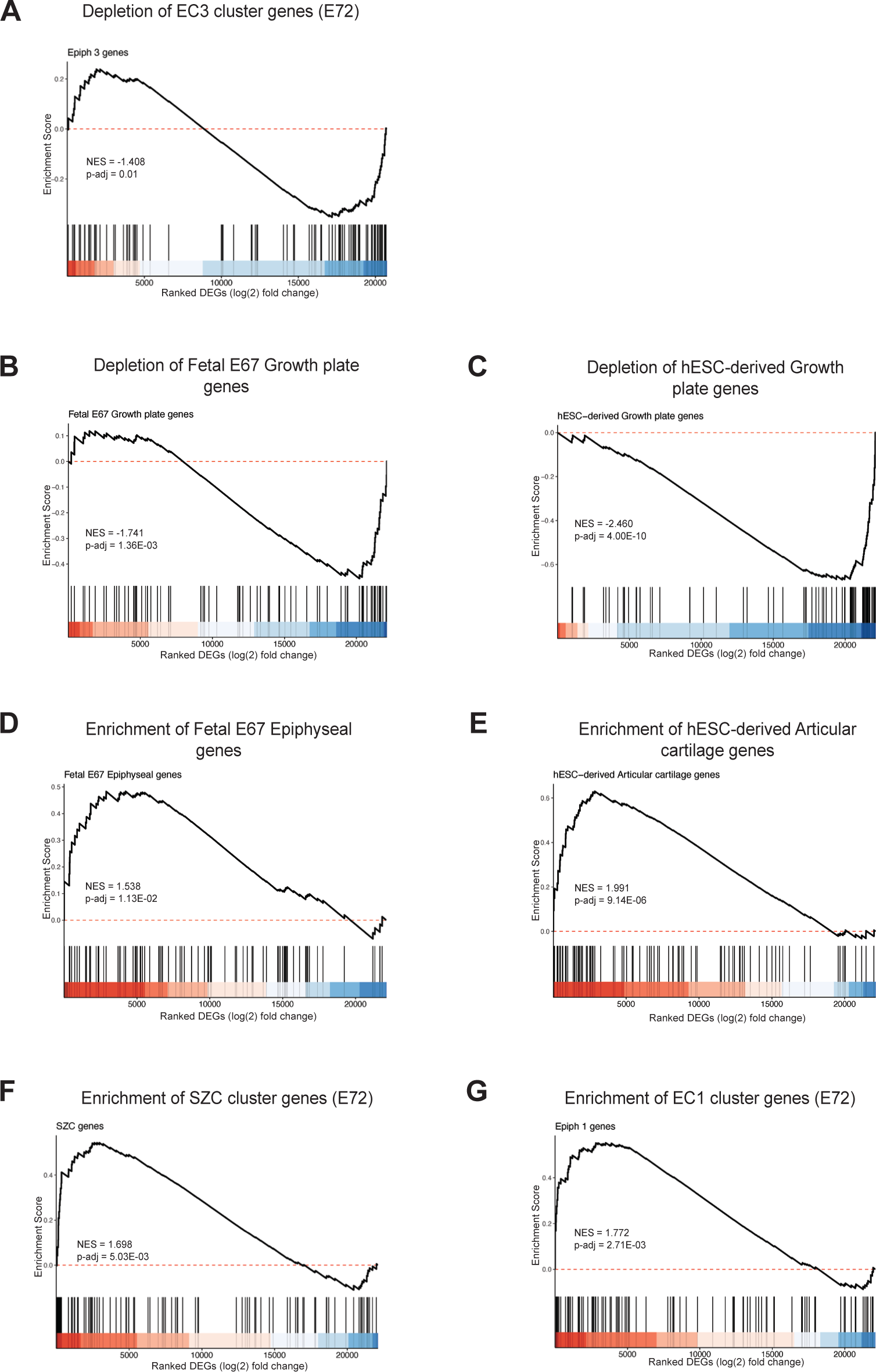
GSEA of preranked genes from indicated reference genesets in NFATC2 OE tissues. (A) Significant depletion of E72 epiphyseal chondrocyte 3 cluster genes in early NFATC2 OE tissues compared to control vehicle treated tissues. (B-C) Significant depletion of genes representing the top DEGs in E67 fetal growth plate cartilage (B) and the top DEGs in hESC-derived growth plate cartilage (C) in mid-stage NFATC2 OE tissues compared to control-vehicle treated tissues. (E-H) Significant enrichment of genes representing the top DEGs in E67 fetal epiphyseal cartilage (E), the top DEGs in hESC-derived articular cartilage (F), the top DEGs in E72 SZC cluster (F), and the top DEGs in E72 epiphyseal chondrocyte 1 cluster (G), in mid-stage NFATC2 OE tissues compared to control-vehicle treated tissues. NES, Normalized GSEA enrichment score. p-adj, adjusted p-value using an optimized false discovery rate approach, was calculated by GSEA. A positive NES indicates enrichment in the NFATC2 OE tissues compared to control tissues, and a negative value indicates a depletion in the NFATC2 OE tissues.

## Supplemental Tables List

**Supplemental Table S1**. Differentially expressed genes between all clusters in the E72 distal femur

**Supplemental Table S2.** Differentially expressed genes between named clusters in the E59 distal femur

**Supplemental Table S3.** Differentially expressed genes between named clusters in the E72 distal femur

**Supplemental Table S4.** Differentially accessible chromatin regions between named clusters in the E59 distal femur

**Supplemental Table S5.** Differentially accessible chromatin regions between named clusters in the E72 distal femur

**Supplemental Table S6.** Differentially active transcription factors between named clusters in the E59 distal femur

**Supplemental Table S7.** Differentially active transcription factors between named clusters in the E72 distal femur

**Supplemental Table S8.** List of eRegulons active in the E59 distal femur, with gene targets

**Supplemental Table S9.** List of eRegulons active in the E72 distal femur, with gene targets

**Supplemental Table S10.** RSS scores for all eRegulons per cluster at E59

**Supplemental Table S11.** RSS scores for all eRegulons per cluster at E72

**Supplemental Table S12.** List of DEGs between replated articular and growth plate-like cartilage, and normalized counts for these genes

**Supplemental Table S13.** List of DEGs between early *NFATC2* overexpression and control hESC-derived growth plate-like cartilage tissues, and normalized counts for these genes

**Supplemental Table S14.** List of DEGs between mid *NFATC2* overexpression and control hESC-derived growth plate-like cartilage tissues, and normalized counts for these genes

**Supplemental Table S15.** GSEA against Hallmark pathway terms in early *NFATC2* overexpression versus control tissues

**Supplemental Table S16.** GSEA against Hallmark pathway terms in mid *NFATC2* overexpression versus control tissues

**Supplemental Table S17.** GSEA against KEGG terms in early *NFATC2* overexpression versus control tissues

**Supplemental Table S18.** GSEA against KEGG terms in mid *NFATC2* overexpression versus control tissues

**Supplemental Table S19.** Top 100 differentially expressed genes between *in vitro*-derived growth plate-like and articular cartilage, and between E67 fetal growth plate and epiphyseal cartilage.

**Supplemental Table S20.** Metadata table of bulk-RNA sequencing samples

**Supplemental Table S21.** Table of gnomAD statistics and GWAS hits for TFs controlling eRegulons

## References

1. Rubin, S. et al. Application of 3D MAPs pipeline identifies the morphological sequence chondrocytes undergo and the regulatory role of GDF5 in this process. Nat Commun 12, 5363 (2021).

2. Kronenberg, H. M. Developmental regulation of the growth plate. Nature 423, 332–6 (2003).

3. Decker, R. S. Articular cartilage and joint development from embryogenesis to adulthood. Seminars in Cell & Developmental Biology 62, 50–56 (2017).

4. Delaere, O. & Dhem, A. Prenatal development of the human pelvis and acetabulum. Acta Orthop Belg 65, 255–60 (1999).

5. Andersen, H. Histochemical studies on the histogenesis of the knee joint and superior tibio-fibular joint in human foetuses. Acta Anat (Basel) 46, 279–303 (1961).

6. Bardeen, C. R. & Lewis, W. H. Development of the limbs, body-wall and back in man. American Journal of Anatomy 1, 1–35 (1901).

7. Bi, W., Deng, J. M., Zhang, Z., Behringer, R. R. & de Crombrugghe, B. Sox9 is required for cartilage formation. Nat Genet 22, 85–9 (1999).

8. Lefebvre, V., Behringer, R. R. & de Crombrugghe, B. L-Sox5, Sox6 and Sox9 control essential steps of the chondrocyte differentiation pathway. Osteoarthritis Cartilage 9 Suppl A, S69-75 (2001).

9. Komori, T. Roles of Runx2 in Skeletal Development. Adv Exp Med Biol 962, 83–93 (2017).

10. Yoshida, C. A. et al. Runx2 and Runx3 are essential for chondrocyte maturation, and Runx2 regulates limb growth through induction of Indian hedgehog. Genes Dev 18, 952–63 (2004).

11. Zhang, C. H. et al. Creb5 coordinates synovial joint formation with the genesis of articular cartilage. Nat Commun 13, 7295 (2022).

12. Wu, M., Wu, S., Chen, W. & Li, Y. P. The roles and regulatory mechanisms of TGF-beta and BMP signaling in bone and cartilage development, homeostasis and disease. Cell Res 34, 101–123 (2024).

13. St-Jacques, B., Hammerschmidt, M. & McMahon, A. P. Indian hedgehog signaling regulates proliferation and differentiation of chondrocytes and is essential for bone formation. Genes Dev 13, 2072–86 (1999).

14. Lauing, K. L. et al. Aggrecan is required for growth plate cytoarchitecture and differentiation. Dev Biol 396, 224–36 (2014).

15. Zhang, B. et al. A human embryonic limb cell atlas resolved in space and time. Nature (2023) doi:10.1038/s41586-023-06806-x.

16. Wu, C. L. et al. Single cell transcriptomic analysis of human pluripotent stem cell chondrogenesis. Nat Commun 12, 362 (2021).

17. Havis, E. et al. Transcriptomic analysis of mouse limb tendon cells during development. Development 141, 3683–96 (2014).

18. Richard, D. et al. Lineage-specific differences and regulatory networks governing human chondrocyte development. Elife 12, (2023).

19. Li, B., Balasubramanian, K., Krakow, D. & Cohn, D. H. Genes uniquely expressed in human growth plate chondrocytes uncover a distinct regulatory network. BMC Genomics 18, 983 (2017).

20. Sun, M. M. & Beier, F. Chondrocyte hypertrophy in skeletal development, growth, and disease. Birth Defects Res C Embryo Today 102, 74–82 (2014).

21. Hao, Y. et al. Integrated analysis of multimodal single-cell data. Cell 184, 3573–3587 e29 (2021).

22. Pei, X. Who is hematopoietic stem cell: CD34+ or CD34-? Int J Hematol 70, 213–5 (1999).

23. Ustanina, S., Carvajal, J., Rigby, P. & Braun, T. The myogenic factor Myf5 supports efficient skeletal muscle regeneration by enabling transient myoblast amplification. Stem Cells 25, 2006–16 (2007).

24. Yang, Z. & Wang, K. K. Glial fibrillary acidic protein: from intermediate filament assembly and gliosis to neurobiomarker. Trends Neurosci 38, 364–74 (2015).

25. Funari, V. A. et al. Cartilage-selective genes identified in genome-scale analysis of non-cartilage and cartilage gene expression. BMC Genomics 8, 165 (2007).

26. Chen, Y. et al. Type-I collagen produced by distinct fibroblast lineages reveals specific function during embryogenesis and Osteogenesis Imperfecta. Nat Commun 12, 7199 (2021).

27. Docheva, D., Hunziker, E. B., Fassler, R. & Brandau, O. Tenomodulin is necessary for tenocyte proliferation and tendon maturation. Mol Cell Biol 25, 699–705 (2005).

28. Schep, A. N., Wu, B., Buenrostro, J. D. & Greenleaf, W. J. chromVAR: inferring transcription-factor-associated accessibility from single-cell epigenomic data. Nat Methods 14, 975–978 (2017).

29. Zhang, C. H. et al. Creb5 establishes the competence for Prg4 expression in articular cartilage. Commun Biol 4, 332 (2021).

30. Zheng, Q. et al. Type X collagen gene regulation by Runx2 contributes directly to its hypertrophic chondrocyte-specific expression in vivo. J Cell Biol 162, 833–42 (2003).

31. Arnold, M. A. et al. MEF2C transcription factor controls chondrocyte hypertrophy and bone development. Dev Cell 12, 377–89 (2007).

32. Bravo Gonzalez-Blas, C., et al. SCENIC+: single-cell multiomic inference of enhancers and gene regulatory networks. Nat Methods 20, 1355–1367 (2023).

33. Craft, A. M. et al. Generation of articular chondrocytes from human pluripotent stem cells. Nat Biotechnol 33, 638–645 (2015).

34. Raftery, R. M., Pregizer, S. K., Kocher, S. & Craft, A. M. Regenerative capacity of human pluripotent stem cell-derived articular chondrocytes in vitro. J Orthop Res (2024) doi:10.1002/jor.25823.

35. Ranger, A. M. et al. The Nuclear Factor of Activated T Cells (Nfat) Transcription Factor Nfatp (Nfatc2) Is a Repressor of Chondrogenesis. Journal of Experimental Medicine 191, 9–22 (2000).

36. Greenblatt, M. B. et al. NFATc1 and NFATc2 repress spontaneous osteoarthritis. Proceedings of the National Academy of Sciences 110, 19914–19919 (2013).

37. Kaminuma, O. et al. Differential contribution of NFATc2 and NFATc1 to TNF-alpha gene expression in T cells. J Immunol 180, 319–26 (2008).

38. Pan, M. G., Xiong, Y. & Chen, F. NFAT gene family in inflammation and cancer. Curr Mol Med 13, 543–54 (2013).

39. Kawane, T. et al. Dlx5 and mef2 regulate a novel runx2 enhancer for osteoblast-specific expression. J Bone Miner Res 29, 1960–9 (2014).

40. Hassan, M. Q. et al. Dlx3 transcriptional regulation of osteoblast differentiation: temporal recruitment of Msx2, Dlx3, and Dlx5 homeodomain proteins to chromatin of the osteocalcin gene. Mol Cell Biol 24, 9248–61 (2004).

41. Chen, C. et al. Mef2a is a positive regulator of Col10a1 gene expression during chondrocyte maturation. Am J Transl Res 15, 4020–4032 (2023).

42. Naya, F. J. et al. Mitochondrial deficiency and cardiac sudden death in mice lacking the MEF2A transcription factor. Nat Med 8, 1303–1309 (2002).

43. Morasso, M. I., Grinberg, A., Robinson, G., Sargent, T. D. & Mahon, K. A. Placental failure in mice lacking the homeobox gene Dlx3. Proc Natl Acad Sci U S A 96, 162–167 (1999).

44. Tachibana, N. et al. RSPO2 defines a distinct undifferentiated progenitor in the tendon/ligament and suppresses ectopic ossification. Sci Adv 8, eabn2138 (2022).

45. Satake, T. et al. Induction of iPSC-derived Prg4-positive cells with characteristics of superficial zone chondrocytes and fibroblast-like synovial cells. BMC Mol Cell Biol 23, 30 (2022).

46. Sugimoto, Y. et al. Scx+/Sox9+ progenitors contribute to the establishment of the junction between cartilage and tendon/ligament. Development 140, 2280–8 (2013).

47. Rodova, M. et al. Nfat1 regulates adult articular chondrocyte function through its age-dependent expression mediated by epigenetic histone methylation. J Bone Miner Res 26, 1974–86 (2011).

48. Zhang, M. et al. Epigenetically mediated spontaneous reduction of NFAT1 expression causes imbalanced metabolic activities of articular chondrocytes in aged mice. Osteoarthritis and Cartilage 24, 1274–1283 (2016).

49. Stuart, T., Srivastava, A., Madad, S., Lareau, C. A. & Satija, R. Single-cell chromatin state analysis with Signac. Nat Methods 18, 1333–1341 (2021).

50. Zhang, Y. et al. Model-based analysis of ChIP-Seq (MACS). Genome Biol 9, R137 (2008).

51. Fornes, O. et al. JASPAR 2020: update of the open-access database of transcription factor binding profiles. Nucleic Acids Res 48, D87–D92 (2020).

52. Meerbrey, K. L. et al. The pINDUCER lentiviral toolkit for inducible RNA interference in vitro and in vivo. Proc Natl Acad Sci U S A 108, 3665–70 (2011).

53. Kanehisa, M. & Goto, S. KEGG: kyoto encyclopedia of genes and genomes. Nucleic Acids Res 28, 27–30 (2000).

54. Liberzon, A. et al. The Molecular Signatures Database (MSigDB) hallmark gene set collection. Cell Syst 1, 417–425 (2015).

55. Subramanian, A. et al. Gene set enrichment analysis: a knowledge-based approach for interpreting genome-wide expression profiles. Proc Natl Acad Sci U S A 102, 15545–50 (2005).

